# Diversification of retinoblastoma protein function associated with cis and trans adaptations

**DOI:** 10.1101/501866

**Authors:** Rima Mouawad, Jaideep Prasad, Dominic Thorley, Pamela Himadewi, Dhruva Kadiyala, Nathan Wilson, Philipp Kapranov, David N. Arnosti

**Author notes:** email addresses for correspondence.

## Abstract

Retinoblastoma proteins are eukaryotic transcriptional co-repressors that play central roles in cell cycle control, among other functions. Although most metazoan genomes encode a single retinoblastoma protein, gene duplications have occurred at least twice: in the vertebrate lineage, leading to three genes encoding Rb, p107, and p130, while separately in the Drosophila lineage an ancestral *Rbf1* gene and a derived *Rbf2* gene. Structurally, Rbf1 resembles p107 and p130 most closely, and mutation of the gene is lethal, while Rbf2 is more divergent, and is not essential for development. Rbf1 has been demonstrated to be a potent repressor of canonical cell-cycle promoters, unlike Rbf2. The retention of *Rbf2* over 60 million years in the entire Drosophila lineage points to essential functions, however. We show here that Rbf2 regulates a broad set of cell growth control related genes, and can antagonize Rbf1 on specific sets of promoters. *Rbf2* null mutants exhibit abnormal development of the female reproductive tract, with reduced egg laying, while heterozygous null mutants exhibit an increased rate of egg deposition, suggesting that the normal function of this protein is critical for optimal control of fertility. The structural alterations found in conserved regions of the *Rbf2* gene suggest that this gene was sub- or neofunctionalized to develop specific regulatory specificity and activity. We define cis regulatory features of Rbf2 target genes that allow preferential repression by this protein, indicating that it is not merely a weaker version of the ancestral protein. The specialization of retinoblastoma function in Drosophila may reflect a parallel evolution found in vertebrates, and raises the possibility that cell growth control is equally important to cell cycle function for this conserved family of transcriptional corepressors.

## Introduction

Retinoblastoma proteins are highly conserved transcriptional corepressors known to be major regulators of cell cycle, differentiation and apoptosis (Burkhart and Sage, 2008). The well-characterized regulation of cell cycle genes involves the binding and inhibition of E2f/DP1 family transcription factors, and subsequent downregulating their target genes, a pivotal role conserved in virtually all multicellular organisms.

The mammalian retinoblastoma family includes three paralogs: Rb, p107 and p130 have overlapping and distinct functions in gene regulation. In humans, germline mutations in *RB1*, the gene for Rb, cause retinoblastomas, and numerous cancers involve somatic mutations in *RB1* or associated pathway genes. Mutations in genes encoding p130 and p107 are less common in tumors, but in an *RB1* mutant background, they modify disease outcomes (Henley and Dick, 2012; Wirt and Sage, 2010). At least eight E2f transcription factors are found in humans and classified into activators (E2f1-3) and repressors (E2f4-8). Rb interacts with E2f1-5. p107 preferentially interacts with E2f4, and p130 with E2f4 and E2f5. The specific interactions of Rb with the activator E2fs may contribute to its distinct cellular functions. Genetic and molecular studies have uncovered specific activities of Rb family proteins in different tissues and cell types, including a role for Rb in senescence (Chicas et al., 2010), and p130 in quiescence (Henley and Dick, 2012), but is it is not fully understood how cellular functions are distributed among the Rb members. Furthermore, the cis regulatory information that leads to preferential association of specific E2f factors and Rb family members is poorly understood.

The presence of three retinoblastoma paralogs in vertebrates is a derived feature, since most metazoans rely on a single retinoblastoma protein to perform cellular functions. The expansion of the retinoblastoma family in vertebrates suggests that the genes may have undergone subfunctionalization and/or neofunctionalization. From a structural point of view, Rb itself is the most derived paralog, as it possesses structural aspects that differ from p107 and p130, which are more similar to an inferred ancestral gene (Wirt and Sage, 2010). The distinct functions acquired by Rb may involve gaining new gene targets related to new functional roles in regulation of apoptosis and differentiation. Interestingly, unlike the gene duplications that impact many other families of transcription factors, retinoblastoma genes tend not to be duplicated in metazoan lineages, with the exception of Drosophila, where a gene duplication ca. 60 million years ago resulted in the expression of two retinoblastoma proteins, Rbf1 and Rbf2, which are found in all characterized genomes of this genus. Thus, Drosophila provides a natural system in which to consider the impact of gene duplication in this important family.

Rbf1 and Rbf2 proteins have similar but not identical expression patterns in early embryogenesis, but in adults, Rbf1 is ubiquitous, whereas Rbf2 is expressed mainly in the ovaries (Keller et al., 2005; Stevaux et al., 2002). Simpler than the vertebrate system, there are two E2f factors in Drosophila, E2f1 which is an activator and E2f2 which is classified as a repressor. Previous work by Dyson and colleagues suggested that Rbf1 interacts with both E2f factors, whereas Rbf2 interacts mainly with E2f2 (Stevaux et al., 2002). These studies showed that when assayed on cell cycle promoters, Rbf2 is a weaker repressor than Rbf1, and few genes are derepressed upon depletion of Rbf2 in cultured S2 cells (Stevaux et al., 2002; Dimova et al., 2003). *Rbf2* null flies do not have a lethal phenotype unlike *Rbf1* null alleles (Stevaux et al., 2005). The conservation of the gene for Rbf2 thus poses a conundrum. Here we explore the activities of Rbf2 and Rbf1 in the context of the intact animal, and show that Rbf2 appears to regulate a large set of genes related to growth control, using unique cis regulatory signals important for specificity. New null alleles of *Rbf2* reveal an important role for Rbf2 in the development and physiological regulation of the ovary. These functions of derived retinoblastoma family members may reflect similar molecular processes that apply to vertebrate paralogs, with application in development and disease.

## Results

### *Rbf2* shows higher divergence than *rbf1* from ancestral lineage, impacting important functional portions of protein sequence

The *Rbf1* and *Rbf2* genes were originally identified by their similarities to mammalian retinoblastoma family genes, including a segment encoding the “pocket” domain critical for interactions with E2f/DP1 (Du et al., 1996; Stevaux et al., 2002). We used multiple sequence alignments to understand conservation of specific segments of these genes within the Drosophila lineage, as well as their relative conservation with other metazoan retinoblastoma genes.

To facilitate our analysis, we divided the protein-coding sequences into three segments: the E2f-binding “pocket” domain (including A and B subdomains), all sequences N-terminal to the pocket, and all sequences C-terminal to the pocket, which include the so-called Instability Element (IE) that is important for stability and activity of pocket proteins (Acharya et al., 2010; Raj et al., 2012; Zhang et al., 2014; Sengupta et al., 2015). Considering Rbf1 protein sequences, the level of conservation for all three domains closely mirrors the overall phylogenetic distances, suggesting that gradual changes in *Rbf1* genes may represent neutral or compensated alterations in the protein. (Figure 1B; Supplementary Figures 1A, 1C, 1D, 1E, Supplementary Table 1A). Rbf2 protein sequences are more divergent overall than Rbf1, especially in sequences of the C-terminus and in the spacer region between the A and B pockets, as well in as the N-terminus (Figure 1A, 1C; Supplementary Figures 1B, 1F, 1G, 1H). Unlike the sequence alignments for Rbf1, Rbf2 sequences can be separated into two clusters; protein sequences from the *melanogaster* subgroup are overall much more similar to each other than those from more distantly related species (*D. ananassae* and others) (Supplementary Figure 1B). The Rbf2 sequences from these more divergent lineages exhibit lower conservation than that observed for Rbf1, meaning that Rbf2 sequences are quite malleable in all Drosophila lineages (Supplementary Table 1A).

**Figure 1:**
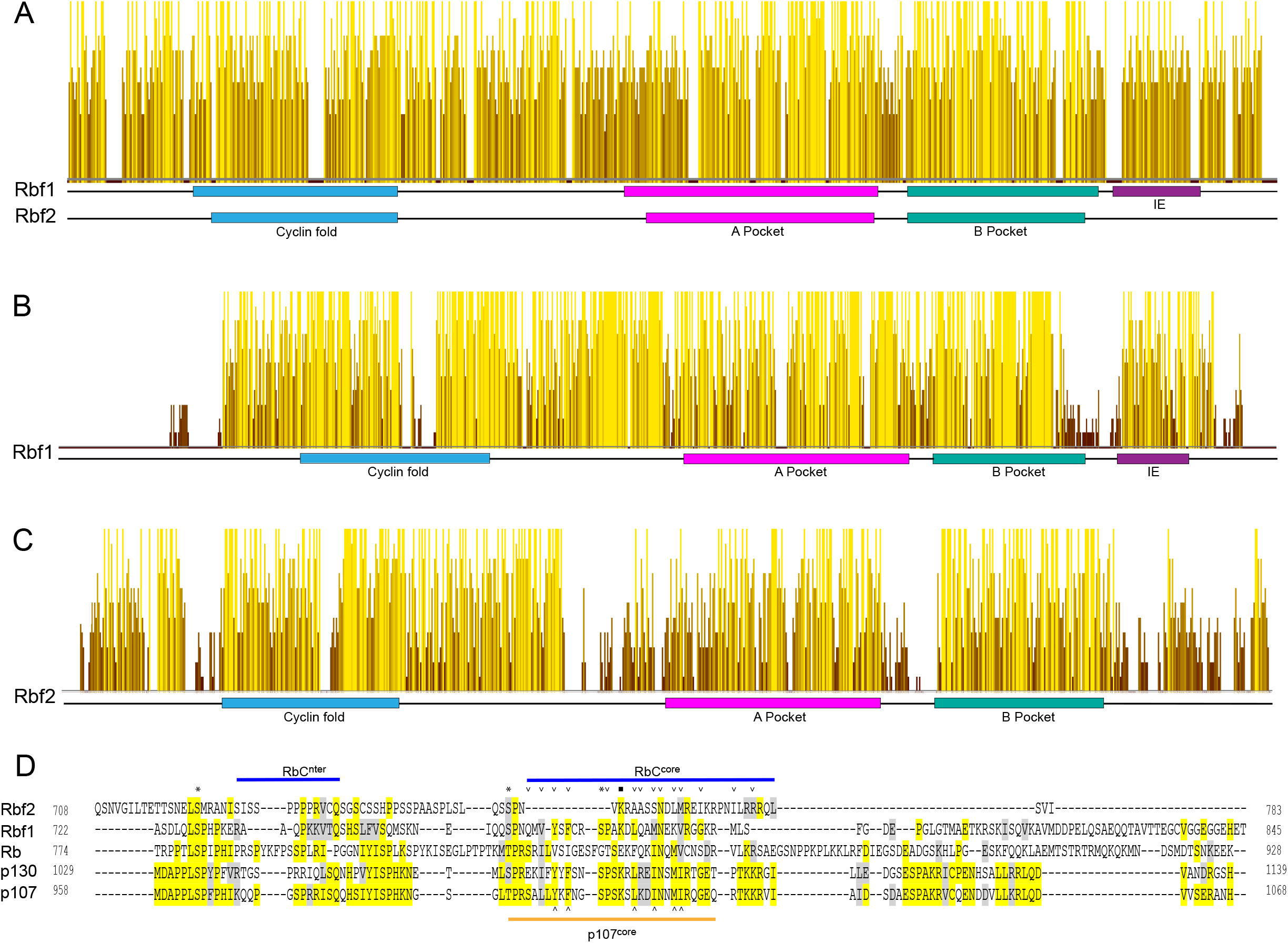
Sequence conservation of retinoblastoma proteins in Drosophila and humans. (A) Pairwise alignment of Drosophila Rbf1 and Rbf2, showing higher conservation in central pocket domains, and lower in C-terminus. (B) Multiple sequence alignment of Rbf1 in 12 Drosophila species; note conservation of C-terminal IE region and A-B pocket spacer. (C) Multiple sequence alignment of Rbf2 in 12 Drosophila species, showing low conservation in C-terminus and in A-B pocket spacer. Height and color of the bars represent percent identity and similarity. Higher yellow bars are more conserved than shorter brown bars. Blue: cyclin fold domain, Pink: A pocket, Green: B pocket, purple: Instability element. (D) Multiple sequence alignment of C-terminus of Drosophila and mammalian retinoblastoma proteins. The yellow color represents conserved residues and grey represents similar residues. Specific portions of the C-terminus involved in direct contacts with E2F/DP1 proteins are highlighted; the RbC^nter^ and RbC^core^ are shown on top of the figure, and the p107^core^ is shown at the bottom. Triangles represent residues that make contacts with E2F/DP marked box domains for both Rb and p107. The asterix denotes conserved serine residues that are targeted for phosphorylation. The K774 residue within the SPAK motif is denoted by a square.

The pocket regions of retinoblastoma proteins are in general the most conserved; within Drosophila, Rbf1 regions A and B show higher conservation among themselves than do the comparable regions of Rbf2 (Supplementary Figure 1A, 1B). Also impacted is the spacer region between the A and B domains: in mammalian p107 and p130 proteins, the spacer regions have unique cyclin/cdk binding and inhibition activity that is absent from Rb, suggesting that changes in this region have functional consequences (Wirt and sage, 2010). In the Drosophila counterparts, the spacer between the A and B pocket domains is well conserved among Drosophila Rbf1 homologs, with a constant length of 19 amino acids (Figure 1B; Supplementary Figure 1D). Rbf2 proteins, in contrast, feature spacer sequences of different lengths and more sequence diversity (Figure 1C; Supplementary Figure 1G).

In the Rbf1 C-terminus, the IE is the most conserved region, which is consistent with our previous studies that this degron is critical for turnover and function (and is also conserved in p107 and p130). Serines 728, 760, and 771 have been shown to mediate regulation by phosphorylation in *D. melanogaster* Rbf1 (Zhang et al., 2014). Lysine 774, which is conserved in p107 and p130, plays an important regulatory role, and is known to be a target of acetylation in the mammalian system (Saeed et al, 2012). These residues are highly conserved in all Rbf1 sequences (Supplementary Figure 1E). Although there are residues of the Rbf2 C-terminus that align with Rbf1 sequences, Rbf2 proteins appear to lack a canonical IE, and only one of the three conserved serine residues found in Rbf1 can be identified (Figure 1D).

Retinoblastoma family proteins contain a cyclin-fold homology domain within the N-terminus. This region is equally well conserved in Rbf1 and Rbf2, with more divergence in Rbf2 sequences in the region between the cyclin fold and the pocket (Figure 1B, 1C). Threonine residue 356 of *D. melanogaster* Rbf1 has been shown to be important for regulation of the protein by phosphorylation, similar to Rb (Burke et al., 2012); this residue is absolutely conserved in Rbf1 proteins; this residue is not conserved in Rbf2 (Supplementary Figure 1C).

The sequence variations for the two gene Drosophila Rbf family may represent a functional interplay between these genes, reflecting neofunctionalization or subfunctionalization. To understand how divergence of Rbf protein sequences in the Drosophila lineage compares with that observed in related arthropod lineages possessing a single *Rbf* gene, we aligned sequences of diverse insect orders, as well as more distantly related chelicerate and crustacean proteins (Supplementary Figure 1I, 1J, 1K). Conserved features noted in Rbf1 are a general feature of homologous proteins; in these genomes, we see a conservation of the N-terminal cyclin fold, the C-terminal IE element and the phosphorylation sites discussed above. Interestingly, the A-B spacer sequence or length is not highly conserved. The diversification in sequence found in Rbf2 proteins in these generally conserved domains points to relaxed constraints on protein structure, perhaps underlying a new cellular role for this protein. We hypothesize that Rbf2 may have diverged faster than Rbf1 because of specialized roles it may have assumed in the physiology and reproduction of different Drosophila species, as indicated by our genetic studies. In this view, Rbf1 may be responsible for regulation of conserved, general functions that are not subject to variation across Drosophila species.

In order to understand how conservation patterns observed for the Drosophila Rbf1 and Rbf2 proteins compare to the other lineage in which this gene family shows duplications, we aligned *D. melanogaster* Rbf1 and Rbf2 with the human Rb, p107, and p130 protein sequences. As previously observed, Rbf1 sequences are more similar to those of p107 and p130 than to those of Rb (Figure 1D). In the C-terminal IE region, specific residues shown to be critical for the selectivity of p107 for E2f4 (Liban et al., 2017) are conserved in Rbf1, but not Rbf2. Interestingly, the human Rb protein is more divergent in this area, suggesting that changes in the IE may be a common mechanism for divergent function of duplicated retinoblastoma protein family members.

### In vivo regulation of embryonic genes by Rbf1 and Rbf2

In previous studies, we mapped in vivo binding profiles of Rbf1 and Rbf2 in the embryo. Rbf2 is found at the promoters of approximately 4,000 genes, while Rbf1 is found at about half that number, in a largely overlapping pattern. The targets of Rbf1 and Rbf2 include ribosomal, cell cycle and signaling genes, however whether these binding events represent direct regulation has not been studied (Acharya et al., 2012, Wei et al., 2015). To determine the effects of Rbf1 and Rbf2 on gene regulation, we induced the expression of each protein using transgenes under the control of a heat shock promoter and performed RNA-seq analysis on 12-18 hour embryos.

After a brief induction of either the Rbf1 or Rbf2 protein, RNA was isolated from embryos, and RNA-seq libraries were prepared for treated or control (heatshock induction with no Rbf transgene) embryos. We filtered the RNA-seq data to focus on genes directly bound by Rbf1 or Rbf2 based on our previous Chip-seq analysis, and removed genes that had low expression levels in all of the samples. We performed unsupervised clustering on the remaining 3937 genes and analyzed five major clusters, with distinct patterns of gene expression across the samples (Figure 2A). Strikingly, Rbf2, which had been characterized as a weak repressor on certain promoters, showed a robust effect on gene expression. All genes in cluster 1 are repressed by Rbf2, with some also exhibiting a weaker repression by Rbf1. On the other hand, Clusters 3 and 5 show a significant upregulation of genes by Rbf2, whereas Rbf1-mediated changes are fewer in number, and weaker in these clusters (Figure 2A; supplementary Table 2A). A number of cell-cycle genes that are repressed by Rbf1 expression in our dataset were found to upregulated in *Rbf1* knock-down cells and *Rbf1* mutant flies (Dimova et al., 2003; Longworth et al., 2012), confirming the physiological relevance of the system used here (Supplementary Table 2B). The dramatically different effects of Rbf1 and Rbf2 expression point to different functions in gene regulation and cellular processes.

**Figure 2:**
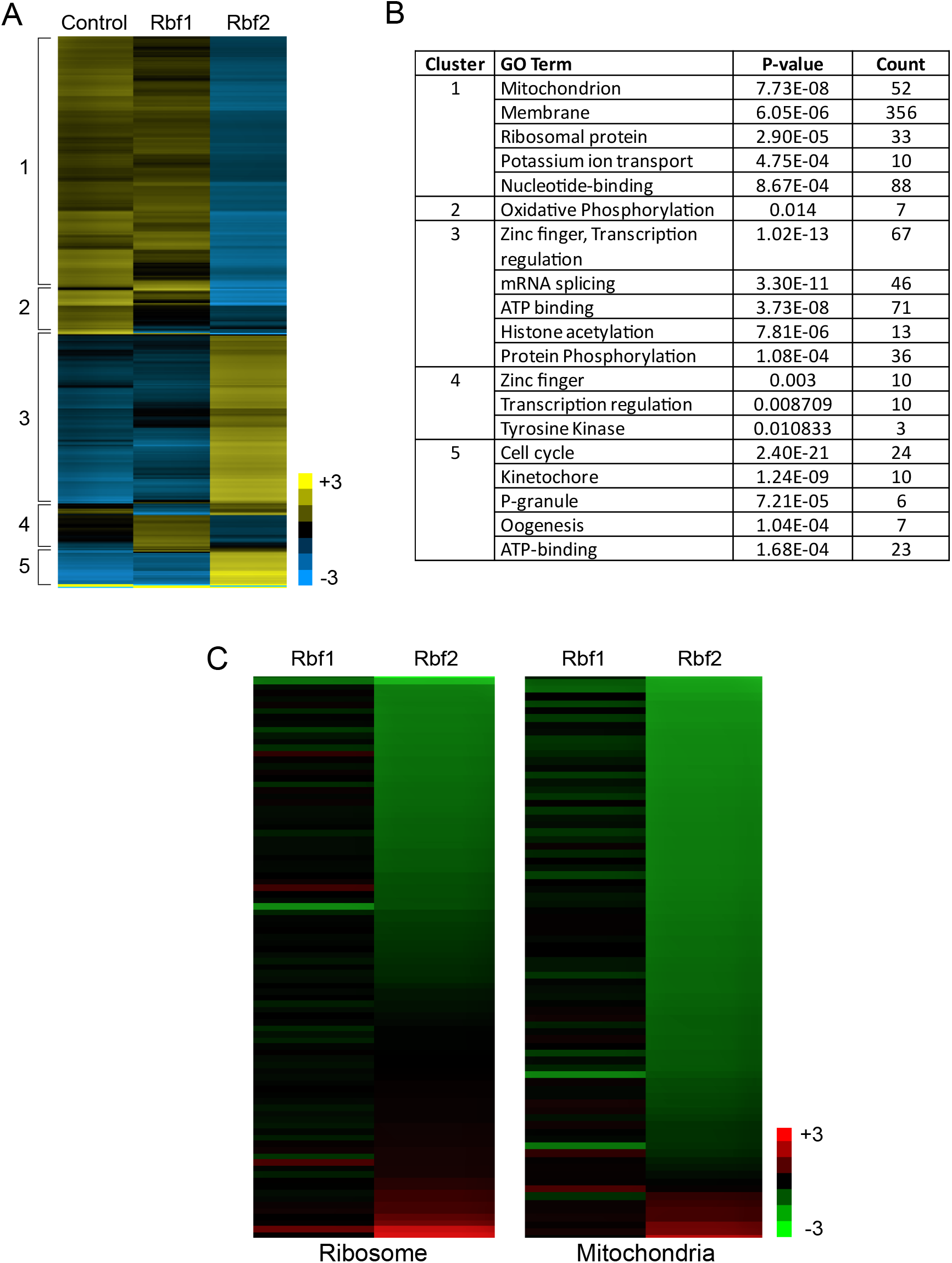
Overexpression of Rbf2 results in profound effects on gene expression in embryos. (A) A heatmap generated by unsupervised clustering of RNA-seq data from Rbf1 and Rbf2 overexpressing embryos, and control embryos. Values represent log transformed RPKM reads and for each gene, the mean is centered to zero. Blue indicates reads below the mean (low expression), black is equal to the mean, and yellow indicates reads above the mean (high expression). Values represent average of three biologic replicates. RPKM < 1 are excluded from the analysis. Only genes bound by Rbf1 or Rbf2 in vivo are included. The heatmap is divided into 5 major clusters based on Euclidean distance. (B) Gene ontology analysis of the 5 clusters based on the DAVID annotation tool. P-values represent significance of enrichment for each category, and the count represents the number of genes in the cluster belonging to each category. (C) Relative gene expression of ribosomal and mitochondrial related genes in Rbf1 or Rbf2 overexpressing embryos, relative to control embryos. Values represent average of three biologic replicates.

To determine the nature of the genes within each cluster, we performed gene ontology analysis using the DAVID annotation tool. Strikingly, among the most enriched categories of Cluster 1 are ribosomal protein and mitochondrial genes, suggesting that Rbf2 may have an important role in control of genes closely linked to cellular growth control (Figure 2B). For specific functional classes of genes, a significant fraction was regulated by Rbf2. For instance, of 93 ribosomal protein genes that are direct targets of Rbf1 or Rbf2, 52 genes show at least 10 *%* repression by Rbf2, and 15 genes by Rbf1. Out of 80 mitochondrial genes that are direct targets, 70 are repressed at least by 10 % by Rbf2 and 23 genes by Rbf1 (Figure 2C). Thus, Rbf2 appears to play a dominant role in regulation of these cell growth-related genes, with Rbf1 playing a secondary role. In the cases where we observe activation by Rbf2, the most enriched categories in Cluster 3 include splicing and transcription regulation, while the Cluster 5 top enriched category is cell cycle. The positive action of Rbf2 overexpression may represent antagonistic action against Rbf1; notably, Cluster 3 and 5 genes have a somewhat higher fraction of promoters co-bound by both Rbf1 and Rbf2 (Supplementary Table 2C).

We considered whether the activation or repression by Rbf2 may relate to the inherent expression levels of targeted genes. Indeed, the majority of genes in Cluster 1 (repressed by Rbf2) were in the top 50% of expression, whereas half of Cluster 5 genes (strongly activated by Rbf2) were in the lowest quartile of expression (Supplementary Figure 2A). Under normal circumstances, the targets in Cluster 5 may be kept inactive by endogenous Rbf1, and competition by Rbf2 upon overexpression may cause them to be derepressed, if Rbf2 is less effective as a repressor. Overall, the functional comparison of Rbf1 and Rbf2 activity in the embryo points to a previously unappreciated role for Rbf2 to regulate a pervasive and functionally distinct set of genes linked to growth regulation.

### 3-Roles for Rbf2 in development and function of the ovary

A previous study generated an *Rbf2* null using deletion of a genomic fragment including the gene, however, this mutation also impacted the neighboring gene, *moira*, and resulted in expression of a fragment of Rbf2 protein (Stevaux et al., 2005). We generated additional *Rbf2* alleles using CRISPR/Cas9, producing four frameshift alleles that truncate the protein N-terminal to the pocket domain, and two in-frame alleles removing five amino acids in two portions of the N-terminus (Figure 3A). Transheterozygous combinations of the null alleles yielded viable flies, consistent with previous reports for viability of the null mutant. Western blot analysis from ovaries of null flies verify the loss of the Rbf2 protein (Figure 3B). Levels of *Rbf2* transcripts are reduced in this background, presumably due to destabilization of the mRNA by translational defects (Figure 4D).

**Figure 3:**
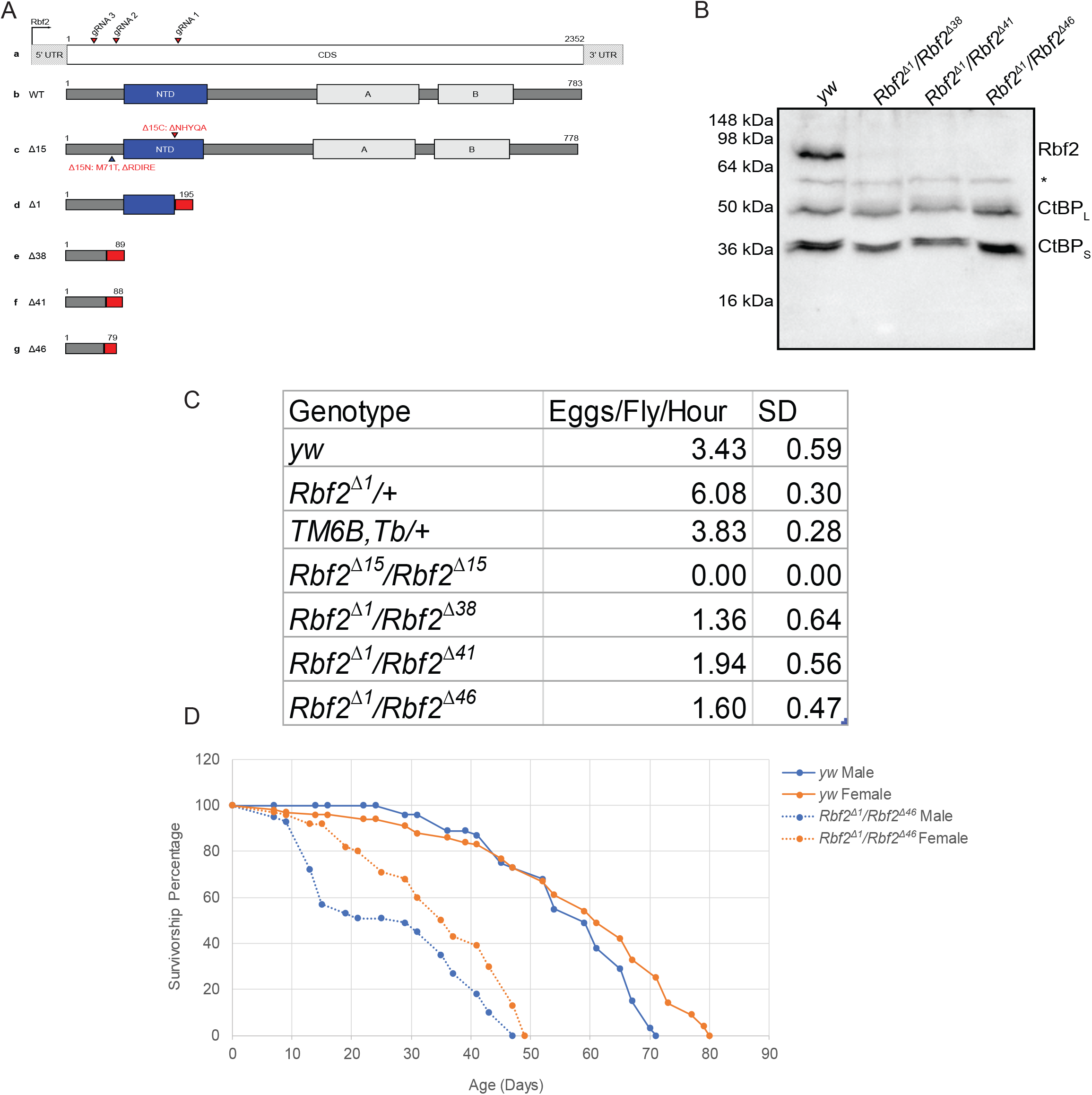
Loss of *Rbf2* affects egg laying and lifespan. (A) Schematic representation of the CRISPR targeting of *Rbf2* and the alleles generated. gRNA1 produced Δ1 and Δ15C alleles, and gRNA2 produced Δ38, Δ41, Δ46 and Δ15N alleles. (B) Western blot indicating loss of Rbf2 protein from *Rbf2* null ovaries from *Rbf2*^△*1*^/*Rbf2*^△*38*^, *Rbf2*^△*1*^/*Rbf2*^△*41*^ and *Rbf2*^△*1*^/*Rbf2*^△*46*^. Anti-CtBP is used as loading control. Ovaries from *yw* flies show the Rbf2 protein. (C) Table showing the number of eggs laid for females of each genotype shown. The numbers represent averages of three biologic replicates and the corresponding standard deviations. (D) Survivorship curve for Rbf2^Δ1^/Rbf2Δ^46^ females and males in comparison to *yw* female and male flies.

**Figure 4:**
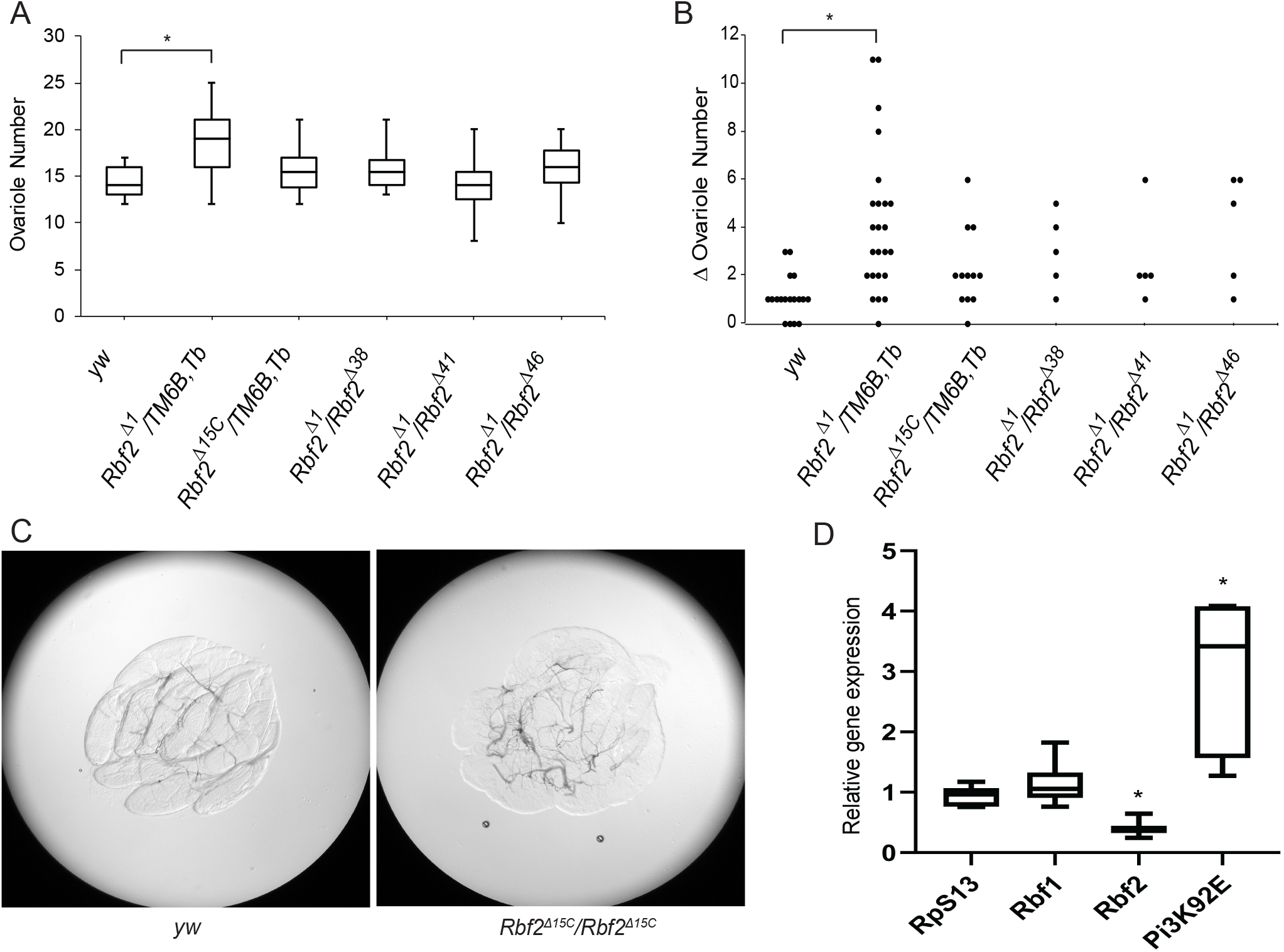
Rbf2 partial loss affects ovariole numbers and complete loss leads to increase in *Pi3K92E* expression. (A) Ovariole counts of individual adult ovaries and (B) difference in ovariole number between ovaries of each female for the following genotypes: *yw* (n = 36), *Rbf2*^△*1*^/*TM6B,Tb* (n = 48), *Rbf2*^△*15C*^/*TM6B,Tb* (n = 24), *Rbf2*^△*1*^/*Rbf2*^△*38*^ (n = 10), *Rbf2*^△*1*^/*Rbf2*^△*41*^ (n = 10), and *Rbf2*^△*1*^/*Rbf2*^△*46*^ (n = 10). (*) indicates p-value<0.01. (C) Images of ovary from *yw* and *Rbf2*^△*15C*^/*Rbf2*^△*15C*^. Images were taken at 10X magnification. (D) Box plot representing relative gene expression of *Rbf2* and *Pi3K92E* from ovaries of *Rbf2*^△*1*^/*Rbf2*^△*46*^ flies in comparison to control *yw* flies. Data represents six biologic replicates. (*) indicates p-value < 0.05.

Although *Rbf2* mutants are viable, the mutants exhibited a spectrum of effects on ovarian development and function, as well as survival. Heterozygous *Rbf2*^Δ1^ females showed a marked increase in egg laying rates, and morphologically distinct, overall larger ovaries filled with a higher fraction of mature oocytes (Figure 3C, and data not shown). The average number of ovarioles per ovary was also ~30% higher in these mutants (Figure 4A, 4B). We found that heterozygous females with *Rbf2*^38^, *Rbf2*^41^, and *Rbf2*^46^ null mutations exhibited similar increases in egg laying, indicating that the partial loss of Rbf2 protein increases egg deposition under these laboratory conditions (data not shown). In contrast to the heterozygotes, homozygous null mutants (*Rbf2*^Δ1^/*Rbf2*^Δ38^, *Rbf2*^Δ1^/*Rbf2*^Δ41^, *Rbf2*^Δ1^/*Rbf2*^Δ46^) laid significantly fewer eggs than wild-type flies (Figure 3C). The lifespan of these flies was significantly shorter than control *yw* flies (Figure 3D showing data for *Rbf2*^Δ1^/*Rbf2*^Δ46^). A number of females completely lacking Rbf2 protein exhibited abnormal morphology of the ovaries, including distorted oocyte shapes, and blackened or greenish oocytes (data not shown). Typically, this was noted on a unilateral basis; both ovaries were not equally affected.

The *Rbf2*^Δ15C^/*Rbf2*^Δ15C^ homozygotes bearing an in-frame deletion of five amino acids in the cyclin fold motif within the N-terminus of the protein were female sterile. The ovaries were very small, with distorted morphology, no discernable germarium or ovariole structures (Figure 4C). Male fertility, on the other hand, was unaffected. *Rbf2*^Δ1^/*Rbf2*^Δ15C^ transheterozygote females had no obvious defects, suggesting that if the in-frame deletion creates a neomorphic protein, there must be a dosage threshold for this phenotype to be displayed.

To understand the null phenotypes at a molecular level, we assessed expression from select target genes in ovaries of *Rbf2*^Δ1^/*Rbf2*^Δ46^ transheterozygous null flies. As expected, *Rp49, Rbf1* and *PCNA* are not affected in the *Rbf2* null flies. Interestingly, another Rbf2 direct target gene, *Pi3K92E* is significantly increased in the null flies in comparison to controls (Figure 4D). This gene encodes the catalytic subunit of class I phosphoinositol-3-kinase that is a component of the insulin signaling pathway and is directly linked to organ growth. Regulation of this gene may contribute to the phenotypes observed.

### Specificity of promoter targeting by the Rbf2 corepressor

To identify cis-regulatory elements that may drive the differential gene regulation by Rbf1 and Rbf2, we performed motif analysis on gene promoters in each cluster of the heat map, focusing on regions under Rbf1 or Rbf2 peaks. Certain motifs were enriched only in specific clusters, suggesting that they may represent binding sites for specificity factors that influence the activity of Rbf proteins (Supplementary Figure 3A). Cluster 1 possessed a motif with similarities to a cell cycle homology region (CHR) motif, and Cluster 3 was specifically enriched in four motifs for known transcription factors, including the Aef1 repressor protein. Cluster-specific motifs were also noted for Clusters 4 and 5.

The E2f motif is uniformly distributed across all clusters, consistent with its important role in mediating E2f/DP1 binding, critical for Rbf recruitment. E2f binding may therefore not be a discriminant for differential Rbf1 and Rbf2. However, the E2f motif is bound by both E2f1 and E2f2; differential binding of E2f1 and E2f2 may affect regulation by Rbf1 and Rbf2. We referred to E2f1 and E2f2 ChIP datasets (Korenjak et al., 2012), and found that percentage of genes bound by E2f2 are somewhat higher in clusters 3 and 5, while E2f1 bound promoters comprise only a small fraction of each cluster (Supplementary Table 3A). Proteins of the Muv/Myb-dREAM complex are known to co-bind promoters with Rb proteins; we note that ChIP data for these proteins (Georlette et al., 2007) identifies a higher fraction of genes in Clusters 3 and 5 (Supplementary Table 3A). These results indicate a potential role for E2f2 and the dREAM complexes, but there does not appear to be a simple “code” for differential regulation by Rbf1 and Rbf2.

To understand the effect of promoter structures on Rbf1 and Rbf2 repression, we assayed the activities of promoters from different classes of genes regulated by the corepressors, using luciferase reporters (Figure 5A). We tested the effects of expression of Rbf1 or Rbf2 on these promoters in S2 cells. The *PCNA* luciferase reporter is strongly repressed by Rbf1, while Rbf2 has no effect on this gene. In contrast, the *CycB* promoter is preferentially repressed by Rbf2, showing a weaker response to Rbf1 overexpression (Figure 5B). We hypothesized that differences between the *PCNA* and *CycB* promoters may involve different interactions by E2f1 and E2f2, therefore, we expressed E2f1 or E2f2 along with these reporters. *PCNA* is robustly induced by E2f1, but there is little or no effect with E2f2 expression. Strikingly, *CycB* is significantly repressed by E2f1, but induced by E2f2 (Figure 5C).

**Figure 5:**
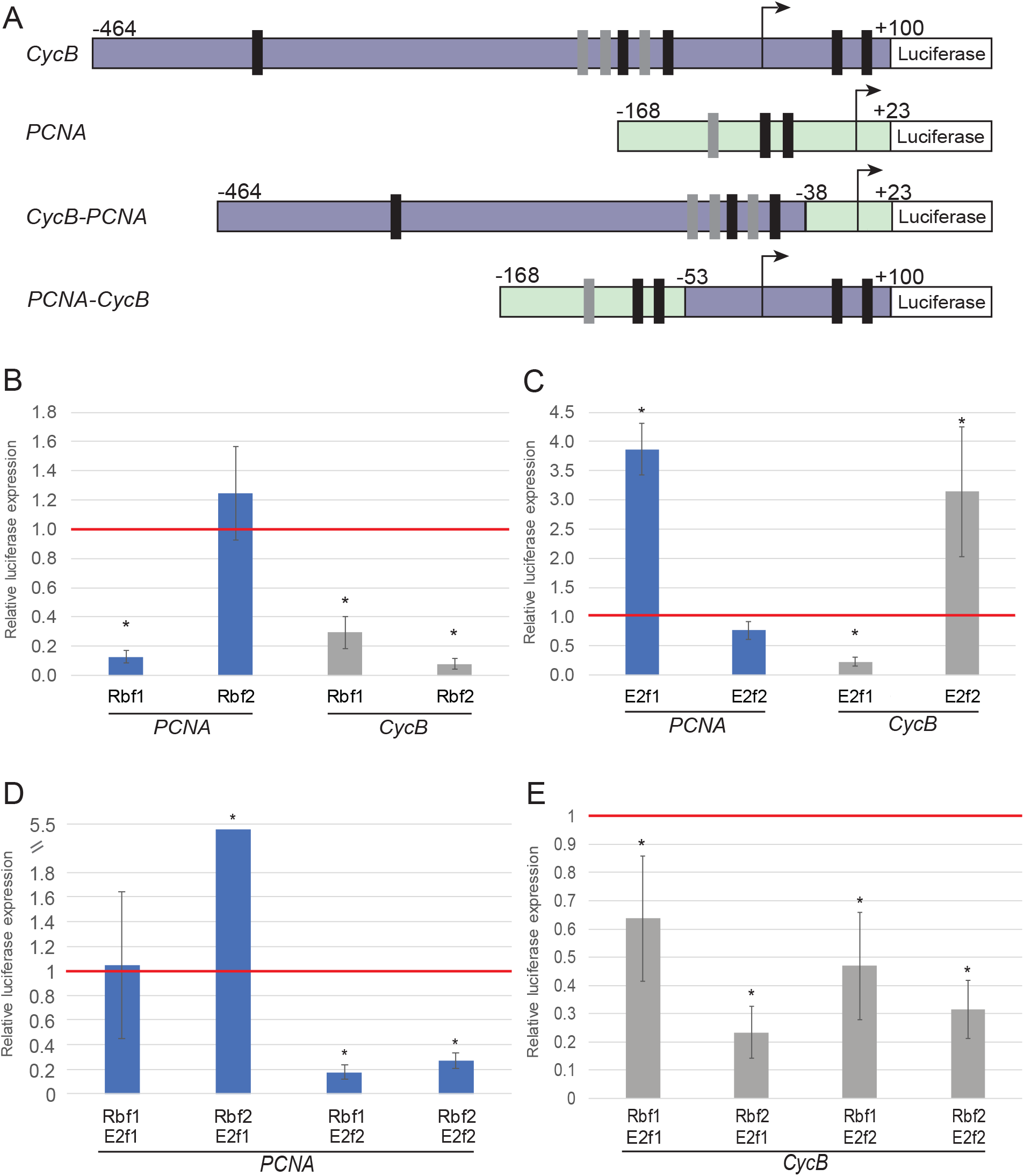
Specific regulation of *PCNA* and *CycB* by Rbf1 and Rbf2. (A) Schematic representation of *CycB, PCNA*, chimeric *CycB-PCNA* and *PCNA-CycB* luciferase reporter genes. (B) Regulation of *PCNA* and *CycB* by Rbf1 and Rbf2. (C) Regulation of *PCNA* and *CycB* by E2F1 and E2F2. (D, E) Combined action of Rbf and E2F proteins on *PCNA* and *CycB* promoters. Luciferase measurements are normalized to expression of the reporters in cells cotransfected with the empty expression vector (no Rbf or E2F gene). Values represent at least three biologic replicates and error bars represent standard deviations. (*) indicates p-value < 0.05.

E2f1 induction of *PCNA* was reversed by co-expression of Rbf1 but not by Rbf2. In contrast, the weak or nonexistent E2f2 repression was substantially enhanced by either Rbf1 or Rbf2 co-expression (Figure 5D). On the promoter that was specifically sensitive to Rbf2, E2f2 induction of *CycB* was reversed by co-expression of Rbf1 or Rbf2 (Figure 5E). This result indicates that on the *CycB* promoter, E2f2 alone does not act as a repressor unless it is bound by Rbf1 or Rbf2. We propose that on this promoter, E2f2 may compete with endogenous E2f1/Rbf complexes, leading to upregulation. Only when Rbf1 or Rbf2 are expressed at higher levels does the E2f2 protein become complexed with a corepressor, and form a repressor complex on the *CycB* promoter. In contrast to E2f2, E2f1 would always be recruited to the promoter complexed with Rbf proteins. These data indicate that E2f1 and E2f2 have different impacts on expression of the *PCNA* and *CycB* promoters, and that the E2f activities are differentially regulated by Rbf1 and Rbf2.

The differential responses of these promoters to E2f and Rbf proteins is undoubtedly mediated by the distinct sequences of these compact promoters. In order to understand the role of the core promoter region of *PCNA* and *CycB*, we created two chimeric reporters (Figure 5A). The first reporter (*CycB-PCNA*) includes *CycB* 5’ sequences (−464 to −53) fused to *PCNA* core promoter region (−38 to +23). A complementary reporter, *PCNA-CycB*, includes *PCNA* 5’ promoter region (−168 to −38) fused to the *CycB* core promoter region (−53 to +100). Introducing the *CycB* core promoter on the *PCNA* reporter (*PCNA-CycB*) permitted repression by Rbf2, although not as strong as for the wild-type *CycB* construct (Figure 6A). Rbf1 repression of this fusion gene was less effective than for the wild-type *PCNA* reporter. Introduction of the *PCNA* core promoter into the *CycB* gene (*CycB-PCNA*) virtually eliminated the strong Rbf2 response; this gene also had weak response to Rbf1 expression (Figure 6A). The *CycB* core promoter appears to play a dominant role in sensitivity to E2f1 and E2f2 expression as well; insertion into the *PCNA* gene turns an E2f1-activated gene into an E2f1 repressed gene, while replacement of this core promoter in *CycB* with the corresponding *PCNA* sequences leads to loss of E2f1 repression, and loss of E2f2 activation (Figure 6B). On *CycB-PCNA*, co-expression of Rbf1 or Rbf2 along with E2f1 or E2f2 showed a response similar to the expression of the E2f proteins alone (Figure 6C). The repression of E2f1 on *PCNA-CycB* is weakened when Rbf1 is coexpressed, while Rbf2 had no impact (Figure 6D). The E2f2 induction of *PCNA-CycB* is reversed after coexpression of either Rbf1 or Rbf2 (Figure 6D).

**Figure 6:**
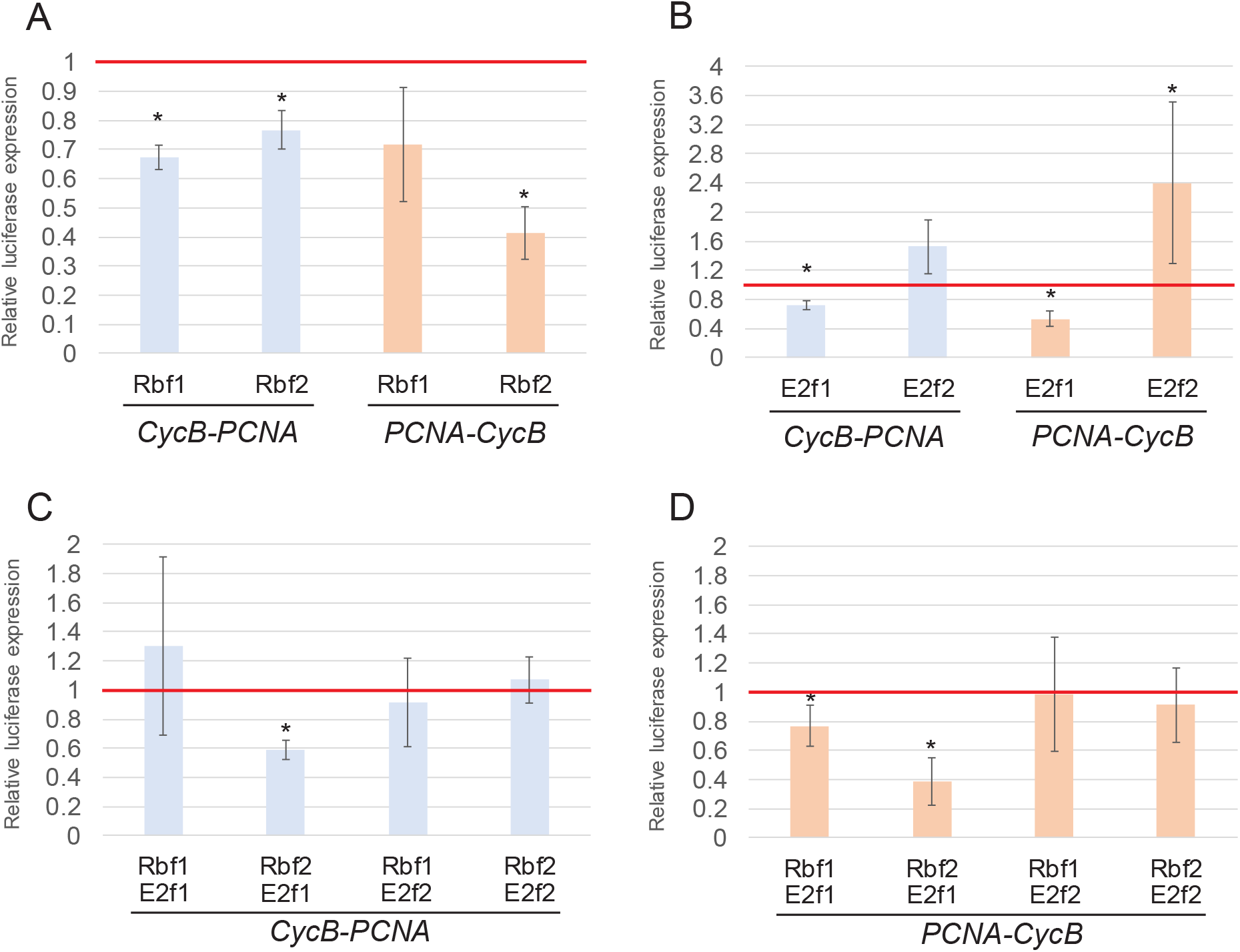
The *CycB* core promoter drives responsiveness to Rbf2. (A) Luciferase reporter assays of chimeric reporters *CycB-PCNA* and *PCNA-CycB* in response to expression of Rbf1 or Rbf2 (B) Effect of expression of E2F1 or E2F2 on chimeric reporters. (C, D) Expression of *CycB-PCNA* or *PCNA-CycB* in response to co-expression of Rbf and E2F proteins. Luciferase measurements are normalized to expression of the reporters in cells cotransfected with the empty expression vector (no Rbf or E2F gene). Values represent at least three biologic replicates and error bars represent standard deviations. (*) indicates p-value < 0.05.

Together, these results indicate that the core promoter region of *CycB* is necessary and sufficient to drive Rbf2 responsiveness, although optimal repression is noted with the endogenous *CycB* regulatory region. The presence of E2f motifs in the *CycB* core promoter suggests that the position or specific sequence of these elements may play a role in differential regulation by the Rbf proteins.

## Discussion

Retinoblastoma protein function appears to be indispensable in almost all eukaryotes, however duplication of retinoblastoma genes has only occurred in selected lineages, including in vertebrates and separately in Drosophila. Whether this duplication involves subfunctionalization, neofunctionalization, or both is not currently understood, but our studies of the derived Rbf2 retinoblastoma protein in Drosophila has uncovered features of unique gene targeting, likely linked to rapid evolutionary changes in several conserved parts of the ancestral protein, as well as connection with fertility that may explain why this gene duplication became locked into Drosophila genomes of diverse species. Although the null alleles we generated confirm the earlier finding by Dyson that *Rbf2* is not strictly required for viability, the impacts on lifespan indicate that in fact on an evolutionary scale, the gene is indispensable.

We speculate that *Rbf2* genes have evolved more rapidly within Drosophila due to specialized functions that are specific to each species. For example, the exact fashion in which transcriptional control is exerted over cell growth-related genes (ribosomal, mitochondrial functions) in response to nutritional signaling may impact the degree to which reproductive strategies are tied to immediate nutritional signals. On the other hand, Rbf1, the major regulator of cell cycle genes, may be more conserved within Drosophila, and more widely in metazoa, because of its role in maintaining core cell cycle functions.

Considering functional domains of retinoblastoma proteins, we find parallel changes in mammals and Drosophila. The C-terminus of retinoblastoma proteins is critical for specific binding to E2f transcription factors. Residues in this domain in the mammalian Rb protein permit specific interaction with E2f1, while limiting p107 and p130 to interactions with E2f4-5 (Rubin et al., 2005; Liban et al., 2017). The C-terminal instability element (IE) region is conserved in the fly Rbf1 as well as the mammalian p107 and p130 proteins; conserved residues in p107 permit specific interaction with E2f4. Strikingly, these residues are conserved in all Drosophila Rbf1 proteins and most arthropods that have a single retinoblastoma protein. The mammalian Rb is divergent in this region; changes in some of the residues allow it to uniquely interact with E2f1 and thus perform Rb-specific functions. Interestingly, Rbf2 is also divergent in this region, perhaps allowing Rbf2 to similarly develop distinct promoter targeting. Indeed, Rbf2 is found at twice the number of promoters as Rbf1, indicating that the binding functions of Rbf1 and Rbf2 are non-identical. Another functional region in retinoblastoma proteins is the spacer region located between the A and B subdomains of the pocket. In mammalian p107 and p130 proteins, the spacer possesses a unique cyclin/cdk binding and inhibition activity that is absent from Rb (Wirt and Sage, 2010). Interestingly, the spacer between the A and B pocket domains is well conserved in Rbf1 among Drosophila, whereas in Rbf2 it is not, possibly affecting the specialized functions of Rbf2 in Drosophila species.

Previous studies of Rbf1 and Rbf2 function focused on these proteins’ activities on reporter genes assayed in cultured cells. On specific cell cycle promoters, Rbf2 has only weak effects compared to Rbf1. Using the embryo as a setting for functional tests of Rbf1 and Rbf2, we found that rather than just being a redundant, and less potent version of Rbf1, Rbf2 has unique effect on distinct classes of genes, such as ribosomal and mitochondrial genes in Cluster I, most of which are directly bound by Rbf2. Interestingly, these are genes that are widely expressed and are viewed as “housekeeping” in nature, however, this designation can obscure the dynamic transcriptional regulation that these genes also undergo. It appears that Rbf2 interactions with these genes are geared to effects that are moderate in nature, changing overall output less than twofold in many cases, a regulation that we deem “soft” repression. Unlike cell cycle target genes that may exhibit complete on/off cycles, these cellular growth-related targets are continuously up and down regulated within specific parameters. Such cybernetic regulation is likely the explanation for the complex transcriptional circuitry found on some Rbf targets, such as the insulin receptor gene, a widely expressed, critical signaling node that includes transcriptional input from Rbf proteins in addition to a dozen additional genetic elements (Wei et al., 2016). Interestingly, the deployment of specific retinoblastoma proteins to cell growth related genes may be a feature that relates to subfunctionalization of these genes (Figure 7C); in human cells, the p130 protein is targeted to many ribosomal protein genes, although the functional relevance remains to be tested.

**Figure 7:**
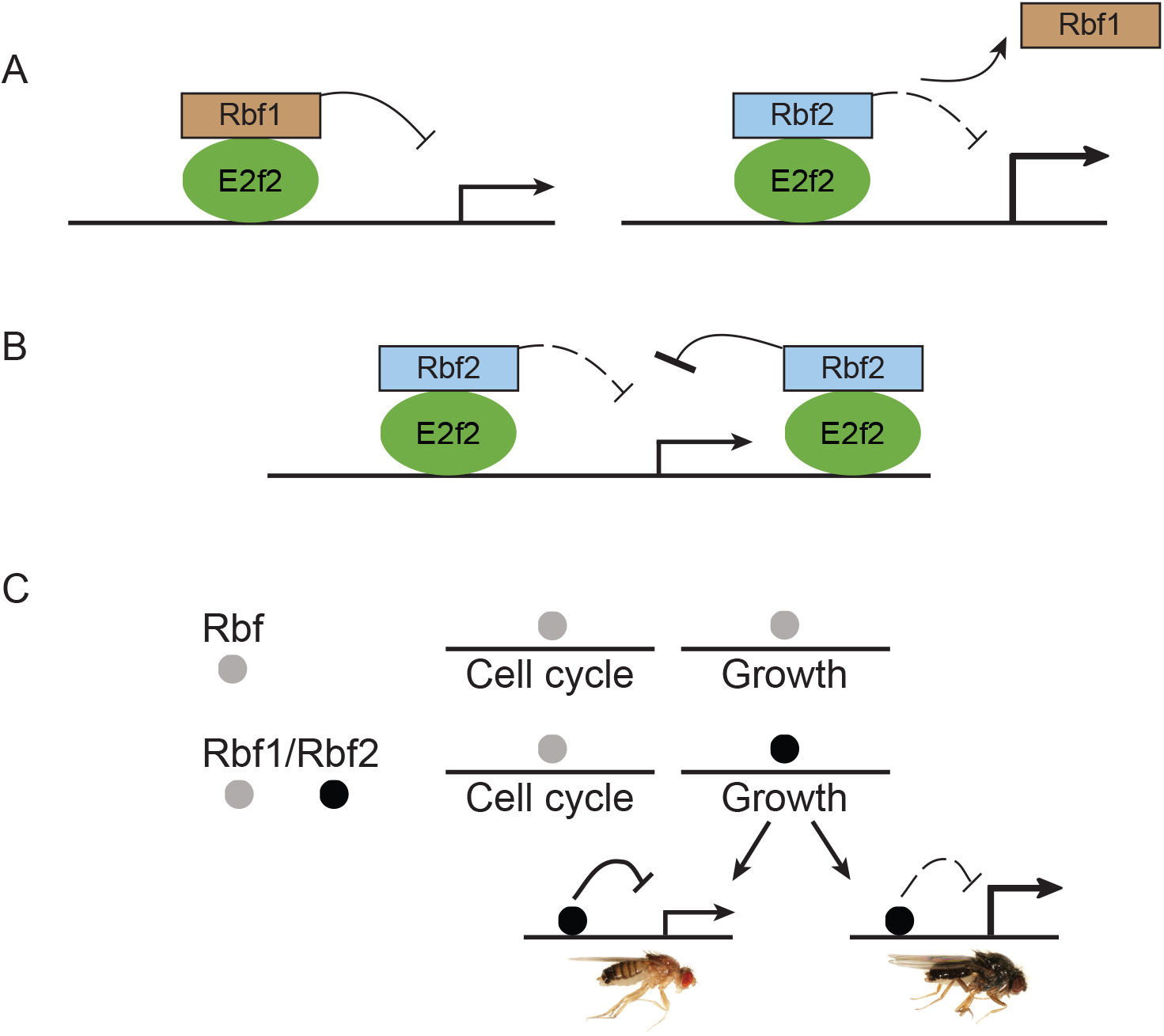
Model for evolved functions of Rbf proteins. (A) Rbf2 competes with Rbf1 binding on E2F2 regulated genes, leading to derepression and optimal gene regulation on certain classes of genes. (B) Rbf2 represses genes more efficiently when it is targeted to the core promoter (as seen in *CycB*). (C) Subfunctionalization of Rbf proteins into cell cycle and cell growth control. Ancestral Rbf proteins regulate cell cycle and cell growth. Evolution of Rbf proteins resulted in subfunctionalization where the more derived Rbf2 protein assumes regulation of cell growth. Rbf2 protein evolved rapidly within Drosophila species to provide optimal growth control and fecundity.

Regarding the biochemistry of transcriptional regulation, numerous studies have pointed to engagement of mammalian retinoblastoma proteins with a wide spectrum of effectors and targets, including E2f proteins, the basal transcriptional machinery and chromatin regulators (Dick, 2007; Fiorentino et al., 2013; Ross et al., 1999). Similar pathways are likely to be invoked in Drosophila, although this area remains to be explored. It is possible that with divergence of Rbf1 and Rbf2, regulatory mechanisms may also differ, with intrinsic differences in the ability to target of basal machinery and recruit histone modifying activities. Alternatively, the finding that Rbf2 regulatory effects appear to be less dramatic than that of Rbf1 may be a function of Rbf2 binding to highly active promoters that are not prone to complete silencing. We explored in depth one instance where Rbf2 exhibits potent repression activity, similar to that found for Rbf1 on its target genes. The ability of Rbf2 to selectively and potently inhibit *CycB* reporter appears to be linked to the unique core promoter sequences, which include putative E2f binding sites. The preferential inhibition by Rbf2 is translated over to chimeric reporters, allowing a switch of specificity from Rbf1 to Rbf2. Such activity may point to a preferential interaction with components of the basal transcription machinery (Figure 7B); indeed, the mammalian Rb protein has been shown to interfere with the basal transcription machinery to regulate E2f target genes (Ross et al., 1999).

Preferential action by Rbf1 or Rbf2 may also relate to the type of E2f protein binding to the promoter; certain E2f sites can be bound by either activator or repressor E2fs (Araki et al., 2003). *PCNA* (responsive to Rbf1 and not Rbf2) may be predominantly regulated by E2f1, while *CycB* may be predominantly regulated by E2f2, as supported by ChIP-seq studies (Korenjak et al., 2012). Our coexpression experiments indicate that there may be preferential association with these promoters by unbound E2f proteins, or E2f associated with Rbf factors (Figure 5); the biochemical basis for such preferential occupancy remains to be elucidated.

A different aspect of Rbf2 function comes from consideration of gene clusters 3 and 5, in which Rbf2 overexpression actually activates genes. Here, antagonism between different retinoblastoma proteins may provide optimal gene regulation by Rb proteins on certain classes of genes. We found that the most potently induced genes were normally expressed at low levels, possibly silenced by Rbf1. We hypothesize that Rbf2 overexpression allows the protein to compete with endogenous Rbf1 binding, allowing the genes to be upregulated due to weaker inhibition (Figure 7A). This mode of regulation is conceivable, given the normal pattern of expression of Rbf1 and Rbf2. The proteins are co-expressed in many different developmental settings, and it is possible that in addition to its unique roles on certain genes, Rbf2 serves as a moderator of Rbf1 activity through such a competitive mechanism. The alternative occupancy of retinoblastoma target gene promoters by different isoforms is well documented in human cells, for instance, p130 replaces Rb on many promoters in quiescent cells; whether the impact on transcription is equivalent, or whether this poises the genes for alternative regulation is unknown (Chicas et al., 2010).

The developmental role of Rbf2 appears to be tightly, but not uniquely, linked to reproduction. Partial loss of function mutants were especially revealing, in that females were found to lay considerably more eggs, as if a “brake” on biosynthesis were lifted – consistent with Rbf2 negative regulation of cell growth related genes. Interestingly, Dyson and colleagues had previously reported a similar phenotype for a disruption in the *Rbf2* locus that they had engineered; what they assumed was a functional null may in fact have residual activity, as an N-terminal portion of the protein may still be expressed (Stevaux et al., 2005).

Gene expression changes in the ovary point to a role for Rbf2 in signaling pathways connected to regulation of oogenesis; we find that *Pi3K92E* is significantly upregulated in *Rbf2* null female ovaries. Studies have shown that female fertility is influenced by the INR/Pi3K signaling pathway (Pritchett and McCall, 2012; Orme et al., 2006). We propose that Rbf2 may be optimizing oogenesis by regulating the INR signaling pathway through Pi3K. In addition to this role in regulation of signaling molecules in the adult, a role for Rbf2 in female reproduction would also involve the development of the ovary, as *Rbf2* heterozygotes often possess ovaries with an increased number of ovarioles, which are specified in larval development. The roles for retinoblastoma proteins in development of the reproductive system appear to be conserved; the conditional knockout of Rb in female mice leads to progressive infertility (Andreu-Vieyra et al., 2008), suggesting that it would be important to examine the role of Rb in regulating fertility in humans as well. In addition to a clear connection to female reproduction, it appears that Rbf2 expression has additional roles in fly physiology. We observed that both male and female *Rbf2* null mutants had shorter lifespans, suggesting that the expression of Rbf2 in embryo and larva outside of reproductive tissues is likely to have significant consequences in both sexes.

Our study has provided important insights on parallel evolution of retinoblastoma paralogs. Mammalian Rb and Drosophila Rbf2 are the most derived proteins in the retinoblastoma family in their respective lineages, and appear to have acquired indispensable new functions that may represent a process of sub-functionalization or neofunctionalization. The findings that Rbf2 is more subject to evolutionary modifications, has acquired unique gene targeting activities, and may play a role in functional antagonism of the ancestral protein appears to mirror similar processes in mammalian systems. These findings present opportunities to explore how biochemical and physiological activities of these conserved transcriptional corepressors are subject to evolutionary modification, and how diverse retinoblastoma protein functions in humans may be better understood in model systems such as Drosophila.

## Materials and Methods

### Protein sequence alignments

Protein sequences of retinoblastoma genes were obtained from FlyBase and NCBI databases. Multiple sequence alignments were generated with Clustal Omega v1.2.4 and ClustalW v2.1 (European bioinformatics institute, EBI) using default settings and manual adjustments. Output files from ClustalW were visualized with Jalview v2.10.5 (EBI).

### Fly stocks

For expression of proteins in the embryo, FLAG-tagged *Rbf1* and *Rbf2* cDNAs were subcloned from the *pAX* vector (Acharya et al. 2010, Wei et al. 2015) into the *pattB* heatshock vector (Kok et al., 2015) using *Hind*III and *Xba*I restriction sites for *Rbf1*. For *Rbf2*, a bridge oligonucleotide containing BglI and *Not*I sites was cloned between *Hind*III and *Xba*I sites, and *Rbf2* cDNA was inserted into *pattB* using these new restriction sites. The plasmids were injected by Rainbow Transgenics into the 51D site on the second chromosome of *yw* flies to generate the homozygous transgenic lines.

### Measurements of gene expression

#### RT-qPCR

*Rbf1* or *Rbf2* transgenes were induced in 12-18 hour embryos by means of a 20-minute heat shock, floating 35 mm apple juice plates with freshly laid embryos on a covered water bath. After 20, 40 or 60 minutes recovery time, RNA was extracted using Total RNA Kit (OMEGA). Control embryos lacking the heat shock transgene were treated similarly to control for nonspecific heat shock effects. cDNA synthesis was performed on total RNA using high capacity cDNA reverse transcription kit (Applied Biosystems). qPCR analysis was performed using Perfecta SYBR Green Fastmix (Quanta Bio). Three biologic replicates were done for both control and transgenic flies (Supplementary Figure 4). To analyze gene expression in *Rbf2* null mutants, RNA was extracted from ovaries using Trizol followed by cDNA synthesis and qPCR analysis, with six biologic replicates from control flies and the mutants. Sequence of primers used are available upon request.

#### RNA-seq analysis

*Rbf1* or *Rbf2* transgenes were induced with a 20-minute heat shock in 12-18 hour embryos. After 60 minute-recovery, RNA was extracted using Total RNA Kit (OMEGA). Control flies were treated similarly to control for the effect of heat shock on gene expression. Poly-A+ RNAs were purified from the total RNA using Oligotex mRNA Mini kit (Qiagen) and were prepared for the SMS essentially as described previously (Kapranov et al., 2010). Sequencing was performed at the SeqLL, LLC facility (Woburn, MA). The SMS reads were processed basically as described before (Kapranov et al., 2010) and aligned to the DM6 version of the *Drosophila melonagaster* genome using indexDPgenomic aligner (Giladi et al., 2010). Uniquely aligned reads were used to generate RPKM values for each transcript annotated in the RefSeq Genes database of the UCSC Genome browser (http://hgdownload.soe.ucsc.edu/goldenPath/dm6/database/refGene.txt.gz) (Kent et al., 2002). Three biologic replicates were done for each sample. Reads lower than RPKM of 1 were removed, which reduced the number of genes to 12,060. Only genes bound by either Rbf1 or Rbf2 based on ChIP-seq dataset (Wei el al., 2015) were further analyzed, for a total of 3937 genes. Unsupervised clustering was performed using Cluster3.0 software, and the heatmap was visualized using JAVA TreeView v1.1.6r4. The counts were log transformed and mean centered, and filtered at 0.3 SD to remove genes with little variation across samples. The heatmap that includes 2795 genes and we decided to analyze the data at the level of five major clusters. Gene ontology analysis was performed for each cluster using DAVID v6.8. Motif analysis was performed on Rbf2 or Rbf1 peak regions of genes in each cluster using MEME-ChIP v4.12.0. Rbf1 and Rbf2 ChIP peak regions are described previously (Wei et al., 2015). Matched motifs were obtained using Tomtom (MEME Suite). The list of Drosophila ribosomal genes was obtained as previously described (Wei el al., 2015), and the list of mitochondrial genes was obtained from MitoDrome database (Sardiello et al., 2003).

#### Generation of novel *Rbf2* alleles with CRISPR

Genomic *Rbf2* target sites were identified at http://tools.flycrispr.molbio.wisc.edu/targetFinder/ (Gratz et al., 2014). Three sites near the 5’ end of the *Rbf2* coding region were selected. Guide-RNAs targeting *ebony* (gRNA-e) and *Rbf2* (gRNA-1, gRNA-2, and gRNA-3) were inserted in vector *pU6-BbsI-chiRNA* (Addgene plasmid #45946) as described (Gratz et al., 2014). *y[1] M{w[+mC]=nos-Cas9.P}ZH-2A w[*]* embryos were injected with each gRNA along with gRNA-e by BestGene Inc. Fly crosses and screening were accomplished using a co-CRISPR strategy adapted from Kane and colleagues (Kane et al., 2017). Injected adult flies, whose germlines potentially contained mutated *Rbf2* alleles, were crossed to the double balancer stock *w[1118]/Dp(1;Y)y[+]; CyO/Bl[1]; TM2, e/TM6B, e, Tb[1]* (Bloomington Drosophila Stock Center #3704) in the parental F(0) generation. Flies in the F (1) generation were scored for ebony body color and tubby pupal shape. These progeny were then crossed to the third chromosome balancer stock *w[1118]; InR[GC25]/TM6B, e, Tb[1]*. Ebony and tubby phenotypes were again scored in the F(2) generation and the flies were crossed inter se to produce homozygous (non-tubby) and balanced (TM6B,Tb) fly lines. Genomic DNA extraction and PCR-amplification of the target-regions was performed on all homozygote and heterozygote lines. Mutations were confirmed by Sanger sequencing. Introgression of the *Rbf2^Δ1^* null allele into a lab stock *yw* flies was performed over 5 generations.

#### Lifespan and fertility assays

Measurement of the *Rbf2* null flies’ lifespans was done as previously described (Linford et al., 2013). 100 females and 100 males from each genotype were separated, and 10 flies were placed in separate vials and maintained at 25°C. After transferring flies into new vials, dead flies were counted and recorded. Assays for fertility were adapted from Stevaux et al. 2005. For the *rbf2*^Δ*1*^/+heterozygote (generated from F(1) of *Rbf2*^Δ*1*^/*TM6B,Tb* crossed to *yw* to produce *rbf2*^Δ*1*^/+ and *TM6B,Tb*/+ flies) the embryos were counted after crossing about 30 staged virgin females with 30 males in laying bottles. After 30-minute preclearing, 3-hour collections were made to determine the egg-laying rate. For the transheterozygous null mutants, the same procedure was done with approximately 15 females crossed to 15 males. For the infertile homozygous *Rbf2*^Δ*15C*^ lines, individual virgin females and virgin males were crossed with *yw* flies to assess female or male sterility. For the introgressed *rbf2*^Δ*1*^ allele, fertility was measured using single female flies.

#### Analysis of mutant ovaries

Ovaries were dissected from staged females on ice-cold PBS with 0.1% Triton X-100. Whole ovaries were directly mounted in 75% glycerol and imaged on a Leica compound microscope under 10X magnification. Ovarioles were split apart and isolated with microsurgical forceps and a fine needle and their number was recorded.

#### Western Blot Analysis

Ovaries were dissected from staged females and homogenized with a polypropylene pestle in lysis buffer (50 mM Tris, pH 8.0, 150 mM NaCl, 1% Triton X-100). The concentration of the extracts was determined via Bradford protein assay, and 50 μg of protein was run per lane in 10% SDS-PAGE gels. Gels were analyzed by Western blot on PVDF membrane using anti-Rbf2 rabbit antibodies (Keller et al., 2005) that bind to the C-terminal end of the protein. Primary antibodies were diluted 1:5000 in TBST (20 mM Tris-Cl, pH 7.5, 120 mM NaCl, 0.1% Tween-20) with 5% nonfat dry milk, and incubation was done overnight at 4°C. Blots were developed using HRP-conjugated goat anti-rabbit secondary antibodies (1:10,000) (30-minute incubation at room temperature) and SuperSignal West Pico chemiluminescent substrate (Pierce).

#### Expression constructs

The *cycB* promoter region (−464 to +100) was cloned into AscI and *SalI* sites in the pAC2T-luciferase vector (Acharya et al., 2010). The PCNA-luciferase reporter (a gift from the Nick Dyson laboratory) was previously described (Acharya et al., 2010; Yamaguchi et al., 1995). The *pIE-E2F1* and *pIE-E2F2* vectors were a gift of the Maxim Frolov laboratory (Frolov et al., 2001). *PCNA-CycB* and *CycB-PCNA* hybrid constructs were synthesized as Gblock gene fragments by IDT (IDTDNA.com) and cloned into the *pAC2T-luciferase* vector (Wei et al., 2015) using *Asc*I and *Sal*I sites.

#### Luciferase reporter assays

Drosophila SL2 cells were cultured in Schneider’s medium (Gibco) supplied with 10% HI-FBS and penicillin-streptomycin (100 units/mL penicillin and 100 μg/mL streptomycin, Gibco). 1.5 million of cells were transfected using Effectene transfection reagent (Qiagen) with 250 ng each of reporter vector, pAX-*Rbf1* or pAX-*Rbf2*, pAX vector as control, and pRL-CMV *Renilla* luciferase reporter. Co-transfection with 250ng of *pIE-E2F1* or *pIE-E2F2* along with *pAX-Rbf1* or *pAX-Rbf2* was also performed, compared to equal amount (500ng) of *pAX* vector as control. Cells were harvested 72 hours post-transfection, and luciferase assays were conducted as described previously (Wei et al., 2015; Acharya et al., 2010).

## Acknowledgements

We thank J. Ravi, J. Rennhack, R.W. Henry, S. Payankaulam, Y. Wei and members of the Arnosti lab for technical assistance and advice. We thank Nick Dyson for sharing the *PCNA* and *Polα* luciferase reporter genes and Maxim Frolov for providing the pIE-Myc-E2f1 and pIE-Myc-E2f2 constructs. We thank the Bloomington Stock Center for Drosophila lines. This research was supported by NIH grant GM124137 to D.N.A. and BEACON consortium to R.M.

## References

Acharya P, Negre N, Johnston J, Wei Y, White KP, Henry RW, Arnosti DN. Evidence for autoregulation and cell signaling pathway regulation from genome-wide binding of the Drosophila retinoblastoma protein. G3 (Bethesda). 2012 Nov;2(11):1459–72. doi: 10.1534/g3.112.004424. Epub 2012 Nov 1. PubMed PMID: 23173097; PubMed Central PMCID: PMC3484676.

Acharya P, Raj N, Buckley MS, Zhang L, Duperon S, Williams G, Henry RW, Arnosti DN. Paradoxical instability-activity relationship defines a novel regulatory pathway for retinoblastoma proteins. Mol Biol Cell. 2010 Nov 15;21(22):3890–901. doi: 10.1091/mbc.E10-06-0520. Epub 2010 Sep 22. PubMed PMID: 20861300; PubMed Central PMCID: PMC2982090.

Andreu-Vieyra C, Chen R, Matzuk MM. Conditional deletion of the retinoblastoma (Rb) gene in ovarian granulosa cells leads to premature ovarian failure. Mol Endocrinol. 2008 Sep;22(9):2141–61. doi: 10.1210/me.2008-0033. Epub 2008 Jul 3. PubMed PMID: 18599617; PubMed Central PMCID: PMC2631371.

Araki K, Nakajima Y, Eto K, Ikeda MA. Distinct recruitment of E2F family members to specific E2F-binding sites mediates activation and repression of the E2F1 promoter. Oncogene. 2003 Oct 23;22(48):7632–41. PubMed PMID: 14576826.

Burke JR, Hura GL, Rubin SM. Structures of inactive retinoblastoma protein reveal multiple mechanisms for cell cycle control. Genes Dev. 2012 Jun 1;26(11):1156–66. doi: 10.1101/gad.189837.112. Epub 2012 May 8. PubMed PMID: 22569856; PubMed Central PMCID: PMC3371405.

Burkhart DL, Sage J. Cellular mechanisms of tumour suppression by the retinoblastoma gene. Nat Rev Cancer. 2008 Sep;8(9):671–82. doi: 10.1038/nrc2399. Review. PubMed PMID: 18650841.

Chicas A, Wang X, Zhang C, McCurrach M, Zhao Z, Mert O, Dickins RA, Narita M, Zhang M, Lowe SW. Dissecting the unique role of the retinoblastoma tumor suppressor during cellular senescence. Cancer Cell. 2010 Apr 13;17(4):376–87. doi: 10.1016/j.ccr.2010.01.023. PubMed PMID: 20385362; PubMed Central PMCID: PMC2889489.

Dick FA. Structure-function analysis of the retinoblastoma tumor suppressor protein - is the whole a sum of its parts? Cell Div. 2007 Sep 13;2:26. PubMed PMID: 17854503; PubMed Central PMCID: PMC2082274.

Dimova DK, Stevaux O, Frolov MV, Dyson NJ. Cell cycle-dependent and cell cycle-independent control of transcription by the Drosophila E2F/RB pathway. Genes Dev. 2003 Sep 15;17(18):2308–20. PubMed PMID: 12975318; PubMed Central PMCID: PMC196467.

Du W, Vidal M, Xie JE, Dyson N. RBF a novel RB-related gene that regulates E2F activity and interacts with cyclin E in Drosophila. Genes Dev. 1996 May 15;10(10):1206–18. PubMed PMID: 8675008.

Fiorentino FP, Marchesi I, Giordano A. On the role of retinoblastoma family proteins in the establishment and maintenance of the epigenetic landscape. J Cell Physiol. 2013 Feb;228(2):276–84. doi: 10.1002/jcp.24141. Review. PubMed PMID: 22718354.

Frolov MV, Huen DS, Stevaux O, Dimova D, Balczarek-Strang K, Elsdon M, Dyson NJ. Functional antagonism between E2F family members. Genes Dev. 2001 Aug 15;15(16):2146–60. PubMed PMID: 11511545; PubMed Central PMCID: PMC312757.

Georlette D, Ahn S, MacAlpine DM, Cheung E, Lewis PW, Beall EL, Bell SP, Speed T, Manak JR, Botchan MR. Genomic profiling and expression studies reveal both positive and negative activities for the Drosophila Myb MuvB/dREAM complex in proliferating cells. Genes Dev. 2007 Nov 15;21(22):2880–96. Epub 2007 Oct 31. PubMed PMID: 17978103; PubMed Central PMCID: PMC2049191.

Giladi E, Healy J, Myers G, Hart C, Kapranov P, Lipson D, Roels S, Thayer E, Letovsky S. Error tolerant indexing and alignment of short reads with covering template families. J Comput Biol. 2010 Oct;17(10):1397–1411. doi: 10.1089/cmb.2010.0005. PubMed PMID: 20937014.

Gratz SJ, Ukken FP, Rubinstein CD, Thiede G, Donohue LK, Cummings AM, O’Connor-Giles KM. Highly specific and efficient CRISPR/Cas9-catalyzed homology-directed repair in Drosophila. Genetics. 2014 Apr;196(4):961–71. doi: 10.1534/genetics.113.160713. Epub 2014 Jan 29. PubMed PMID: 24478335; PubMed Central PMCID: PMC3982687.

Henley SA, Dick FA. The retinoblastoma family of proteins and their regulatory functions in the mammalian cell division cycle. Cell Div. 2012 Mar 14;7(1):10. doi: 10.1186/1747-1028-7-10. PubMed PMID: 22417103; PubMed Central PMCID: PMC3325851.

Kane NS, Vora M, Varre KJ, Padgett RW. Efficient Screening of CRISPR/Cas9-Induced Events in Drosophila Using a Co-CRISPR Strategy. G3 (Bethesda). 2017 Jan 5;7(1):87–93. doi: 10.1534/g3.116.036723. PubMed PMID: 27793971; PubMed Central PMCID: PMC5217126.

Kapranov P, St Laurent G, Raz T, Ozsolak F, Reynolds CP, Sorensen PH, Reaman G, Milos P, Arceci RJ, Thompson JF, Triche TJ. The majority of total nuclear-encoded non-ribosomal RNA in a human cell is ‘dark matter’ un-annotated RNA. BMC Biol. 2010 Dec 21;8:149. doi: 10.1186/1741-7007-8-149. Erratum in: BMC Biol. 2011;9:86. PubMed PMID: 21176148; PubMed Central PMCID: PMC3022773.

Keller SA, Ullah Z, Buckley MS, Henry RW, Arnosti DN. Distinct developmental expression of Drosophila retinoblastoma factors. Gene Expr Patterns. 2005 Feb;5(3):411–21. PubMed PMID: 15661648.

Kent WJ, Sugnet CW, Furey TS, Roskin KM, Pringle TH, Zahler AM, Haussler D. The human genome browser at UCSC. Genome Res. 2002 Jun;12(6):996–1006. PubMed PMID: 12045153; PubMed Central PMCID: PMC186604.

Kok K, Ay A, Li LM, Arnosti DN. Genome-wide errant targeting by Hairy. Elife. 2015 Aug 25;4. doi: 10.7554/eLife.06394. PubMed PMID: 26305409; PubMed Central PMCID: PMC4547095.

Korenjak M, Anderssen E, Ramaswamy S, Whetstine JR, Dyson NJ. RBF binding to both canonical E2F targets and noncanonical targets depends on functional dE2F/dDP complexes. Mol Cell Biol. 2012 Nov;32(21):4375–87. doi: 10.1128/MCB.00536-12. Epub 2012 Aug 27. PubMed PMID: 22927638; PubMed Central PMCID: PMC3486151.

Liban TJ, Medina EM, Tripathi S, Sengupta S, Henry RW, Buchler NE, Rubin SM. Conservation and divergence of C-terminal domain structure in the retinoblastoma protein family. Proc Natl Acad Sci U S A. 2017 May 9;114(19):4942–4947. doi: 10.1073/pnas.1619170114. Epub 2017 Apr 24. PubMed PMID: 28439018; PubMed Central PMCID: PMC5441720.

Linford NJ, Bilgir C, Ro J, Pletcher SD. Measurement of lifespan in Drosophila melanogaster. J Vis Exp. 2013 Jan 7;(71). pii: 50068. doi: 10.3791/50068. PubMed PMID: 23328955; PubMed Central PMCID: PMC3582515.

Longworth MS, Walker JA, Anderssen E, Moon NS, Gladden A, Heck MM, Ramaswamy S, Dyson NJ. A shared role for RBF1 and dCAP-D3 in the regulation of transcription with consequences for innate immunity. PLoS Genet. 2012;8(4):e1002618. doi: 10.1371/journal.pgen.1002618. Epub 2012 Apr 5. PubMed PMID: 22496667; PubMed Central PMCID: PMC3320600.

Orme MH, Alrubaie S, Bradley GL, Walker CD, Leevers SJ. Input from Ras is required for maximal PI(3)K signalling in Drosophila. Nat Cell Biol. 2006 Nov;8(11):1298–302. Epub 2006 Oct 15. PubMed PMID: 17041587.

Pritchett TL, McCall K. Role of the insulin/Tor signaling network in starvation-induced programmed cell death in Drosophila oogenesis. Cell Death Differ. 2012 Jun;19(6):1069–79. doi: 10.1038/cdd.2011.200. Epub 2012 Jan 13. PubMed PMID: 22240900; PubMed Central PMCID: PMC3354059.

Raj N, Zhang L, Wei Y, Arnosti DN, Henry RW. Rbf1 degron dysfunction enhances cellular DNA replication. Cell Cycle. 2012 Oct 15;11(20):3731–8. doi: 10.4161/cc.21665. Epub 2012 Aug 16. PubMed PMID: 22895052; PubMed Central PMCID: PMC3495815.

Ross JF, Liu X, Dynlacht BD. Mechanism of transcriptional repression of E2F by the retinoblastoma tumor suppressor protein. Mol Cell. 1999 Feb;3(2):195–205. PubMed PMID: 10078202.

Rubin SM, Gall AL, Zheng N, Pavletich NP. Structure of the Rb C-terminal domain bound to E2F1-DP1: a mechanism for phosphorylation-induced E2F release. Cell. 2005 Dec 16;123(6):1093–106. PubMed PMID: 16360038.

Saeed M, Schwarze F, Loidl A, Meraner J, Lechner M, Loidl P. In vitro phosphorylation and acetylation of the murine pocket protein Rb2/p130. PLoS One. 2012;7(9):e46174. doi: 10.1371/journal.pone.0046174. Epub 2012 Sep 24. PubMed PMID: 23029429; PubMed Central PMCID: PMC3454344.

Sardiello M, Licciulli F, Catalano D, Attimonelli M, Caggese C. MitoDrome: a database of Drosophila melanogaster nuclear genes encoding proteins targeted to the mitochondrion. Nucleic Acids Res. 2003 Jan 1;31(1):322–4. PubMed PMID: 12520013; PubMed Central PMCID: PMC165570.

Sengupta S, Lingnurkar R, Carey TS, Pomaville M, Kar P, Feig M, Wilson CA, Knott JG, Arnosti DN, Henry RW. The Evolutionarily Conserved C-terminal Domains in the Mammalian Retinoblastoma Tumor Suppressor Family Serve as Dual Regulators of Protein Stability and Transcriptional Potency. J Biol Chem. 2015 Jun 5;290(23):14462–75. doi: 10.1074/jbc.M114.599993. Epub 2015 Apr 22. PubMed PMID: 25903125; PubMed Central PMCID: PMC4505513.

Stevaux O, Dimova D, Frolov MV, Taylor-Harding B, Morris E, Dyson N. Distinct mechanisms of E2F regulation by Drosophila RBF1 and RBF2. EMBO J. 2002 Sep 16;21(18):4927–37. PubMed PMID: 12234932; PubMed Central PMCID: PMC126297.

Stevaux O, Dimova DK, Ji JY, Moon NS, Frolov MV, Dyson NJ. Retinoblastoma family 2 is required in vivo for the tissue-specific repression of dE2F2 target genes. Cell Cycle. 2005 Sep;4(9):1272–80. Epub 2005 Sep 29. PubMed PMID: 16082225.

Wei Y, Gokhale RH, Sonnenschein A, Montgomery KM, Ingersoll A, Arnosti DN. Complex cis-regulatory landscape of the insulin receptor gene underlies the broad expression of a central signaling regulator. Development. 2016 Oct 1;143(19):3591–3603. PubMed PMID: 27702787; PubMed Central PMCID: PMC5087611.

Wei Y, Mondal SS, Mouawad R, Wilczyński B, Henry RW, Arnosti DN. Genome-Wide Analysis of Drosophila RBf2 Protein Highlights the Diversity of RB Family Targets and Possible Role in Regulation of Ribosome Biosynthesis. G3 (Bethesda). 2015 May 20;5(7):1503–15. doi: 10.1534/g3.115.019166. PubMed PMID: 25999584; PubMed Central PMCID: PMC4502384.

Wirt SE, Sage J. p107 in the public eye: an Rb understudy and more. Cell Div. 2010 Apr 2;5:9. doi: 10.1186/1747-1028-5-9. PubMed PMID: 20359370; PubMed Central PMCID: PMC2861648.

Yamaguchi M, Hayashi Y, Matsukage A. Essential role of E2F recognition sites in regulation of the proliferating cell nuclear antigen gene promoter during Drosophila development. J Biol Chem. 1995 Oct 20;270(42):25159–65. PubMed PMID: 7559650.

Zhang L, Wei Y, Pushel I, Heinze K, Elenbaas J, Henry RW, Arnosti DN. Integrated stability and activity control of the Drosophila Rbf1 retinoblastoma protein. J Biol Chem. 2014 Sep 5;289(36):24863–73. doi: 10.1074/jbc.M114.586818. Epub 2014 Jul 21. PubMed PMID: 25049232; PubMed Central PMCID: PMC4155655.

