## Supplementary figure and table legends for "Diversification of retinoblastoma protein function associated with cis and trans adaptations"

Supplementary figure 1A:

Percent identity of from multiple sequence alignment of Rbf1 from Drosophila species against D. melanogaster for the N-terminus, pocket and C-terminus. Percent identity resulted from multiple sequence alignments using Clustal Omega.

Supplementary figure 1B:

Percent identity of from multiple sequence alignment of Rbf2 from Drosophila species against D. melanogaster for the N-terminus, pocket and C-terminus. Percent identity resulted from multiple sequence alignments using Clustal Omega.

Supplementary figure 1C:

Multiple sequence alignment for Rbf1 N-terminus within Drosophila species. Yellow shade represents conserved residues and grey represents similar residues to D. melanogaster. The Cyclin fold domain is shown as black line. The (*) denotes conserved Thr356 residue.

Supplementary figure 1D:

Multiple sequence alignment for Rbf1 pocket domain within Drosophila species. Yellow shade represents conserved residues and grey represents similar residues to *D. melanogaster*. A and B pocket subdomains are shown marking the 19-residue spacer region.

Supplementary figure 1E:

Multiple sequence alignment for Rbf1 C-terminus within Drosophila species. Yellow shade represents conserved residues and grey represents similar residues to *D. melanogaster*. The IE region is shown as black line. The p107^core^ region is shown as red line. Triangles indicate residues that interact with E2F/DP marked box domain.

Supplementary figure 1F:

Multiple sequence alignment for Rbf2 N-terminus within Drosophila species. Yellow shade represents conserved residues and grey represents similar residues to *D. melanogaster*. The cyclin fold region is shown in black.

Supplementary figure 1G:

Multiple sequence alignment for Rbf2 pocket domain within Drosophila species. Yellow shade represents conserved residues and grey represents similar residues to *D. melanogaster*. A and B pocket subdomains are shown.

Supplementary figure 1H:

Multiple sequence alignment for Rbf2 C-terminus within Drosophila species. Yellow shade represents conserved residues and grey represents similar residues to *D. melanogaster*.

Supplementary figure 1I:

Multiple sequence alignment of N-terminus of D. melanogaster Rbf1 and the following arthropod species: D. plexippus (Lepidoptera), A. cerena (Lepidoptera), T. castaneum (Coleoptera), F. occidentalis (Thysanpotera), M. persicae (Hemiptera), C. secundus (Isoptera), F. candida (Hymenoptera), P. tepidariorum (Spider), and P. vannamei (White-legged Shrimp). The yellow shade represents conserved residues and grey represents similar residues. The cyclin fold of D. melanogaster is shown on top of the figure.

Supplementary figure 1J:

Multiple sequence alignment of D. melanogaster Rbf1 pocket and the following arthropod species: D. plexippus (Lepidoptera), *A. cerena* (Lepidoptera), T. castaneum (Coleoptera), F. occidentalis (Thysanpotera), M. persicae (Hemiptera), C. secundus (Isoptera), F. candida (Hymenoptera), P. tepidariorum (Spider), and P. vannamei (White-legged Shrimp). The yellow shade represents conserved residues and grey represents similar residues. The A and B pocket subdomains are indicated.

Supplementary figure 1K:

Multiple sequence alignment of D. melanogaster Rbf1 C-terminus and the following arthropod species: D. plexippus (Lepidoptera), A. cerena (Lepidoptera), T. castaneum (Coleoptera), F. occidentalis (Thysanpotera), M. persicae (Hemiptera), C. secundus (Isoptera), F. candida (Egonatha), P. tepidariorum (Spider), and P. vannamei (White-legged Shrimp). The yellow shade represents conserved residues and grey represents similar residues. The A and B pocket subdomains are indicated. The IE region is shown as black line. The p107^core^ region is shown as red line. Triangles indicate residues that interact with E2F/DP marked box domain.

Supplementary figure 2A:

Bar graph showing expression levels of genes within each cluster of the heatmap. The expression levels were determined from the RPKM values of the genes in the control samples. The RPKM values were ranked from high to low and divided into quartiles.

Supplementary figure 3A:

Motif analysis of the Rbf1/Rbf2 bound promoter regions of genes within each cluster of the heatmap. The name of the TF to which the motif corresponds to is shown on left of the motif logo.

Supplementary figure 4:

Kinetics of gene expression after induction of Rbf1 protein in 12-18hr embryos. *Rp49* is used as control. Data represents average of three biologic replicates. Error bars indicate standard deviation. (*) indicates p-value <0.05.

Supplementary Table 1A:

Spreadsheet showing percent identity values from multiple sequence alignments of Rbf1 and Rbf2 among Drosophila species. Percent identity values were calculated in Clustal Omega alignment tool.

Supplementary Table 2A:

RNA-seq analysis results showing gene expression changes after induction of Rbf1 or Rbf2 in embryo. The changes represent relative expression in comparison to control embryos. Changes in gene expression of more than 20% are counted as up or down, otherwise no change.

Supplementary Table 2B:

Spreadsheet indicating genes that have opposite effects in Rbf1 overexpression and KD (from Longworth et al and Dimova et al).

Supplementary Table 2C:

Ratio of genes that are shown to be occupied by Rbf1 or Rbf2 or both in previous Chip-seq dataset (Wei et al).

Supplementary Table 3A:

Ratio of genes that are shown to be occupied by E2F1, E2F2 or DREAM complex as shown in previous Chip-seq datasets (Dyson, georlette).
