## Supplementary figures and images for "Diversification of retinoblastoma protein function associated with cis and trans adaptations"

### Supplementary Figure1A

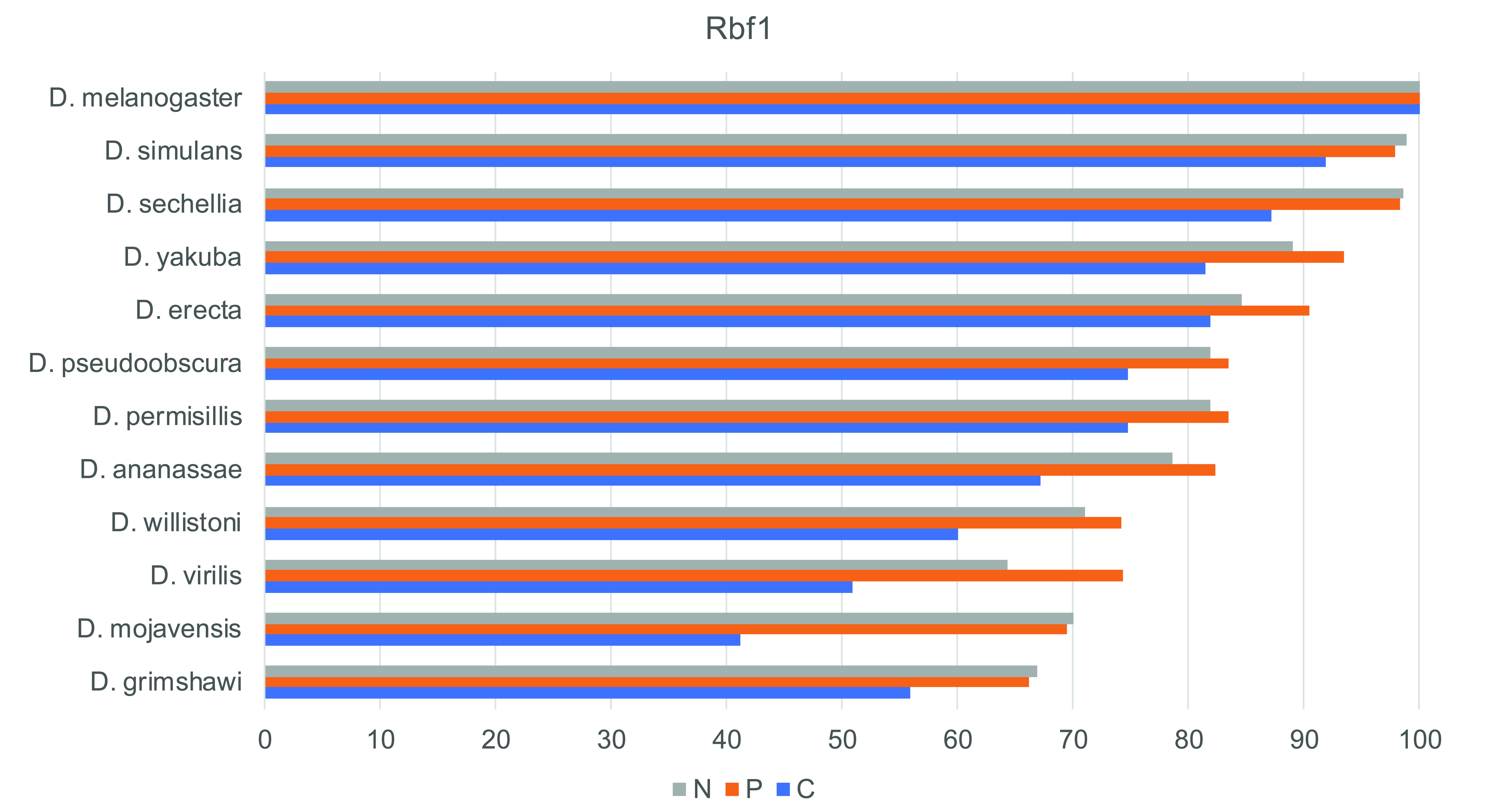

### Supplementary Figure1B

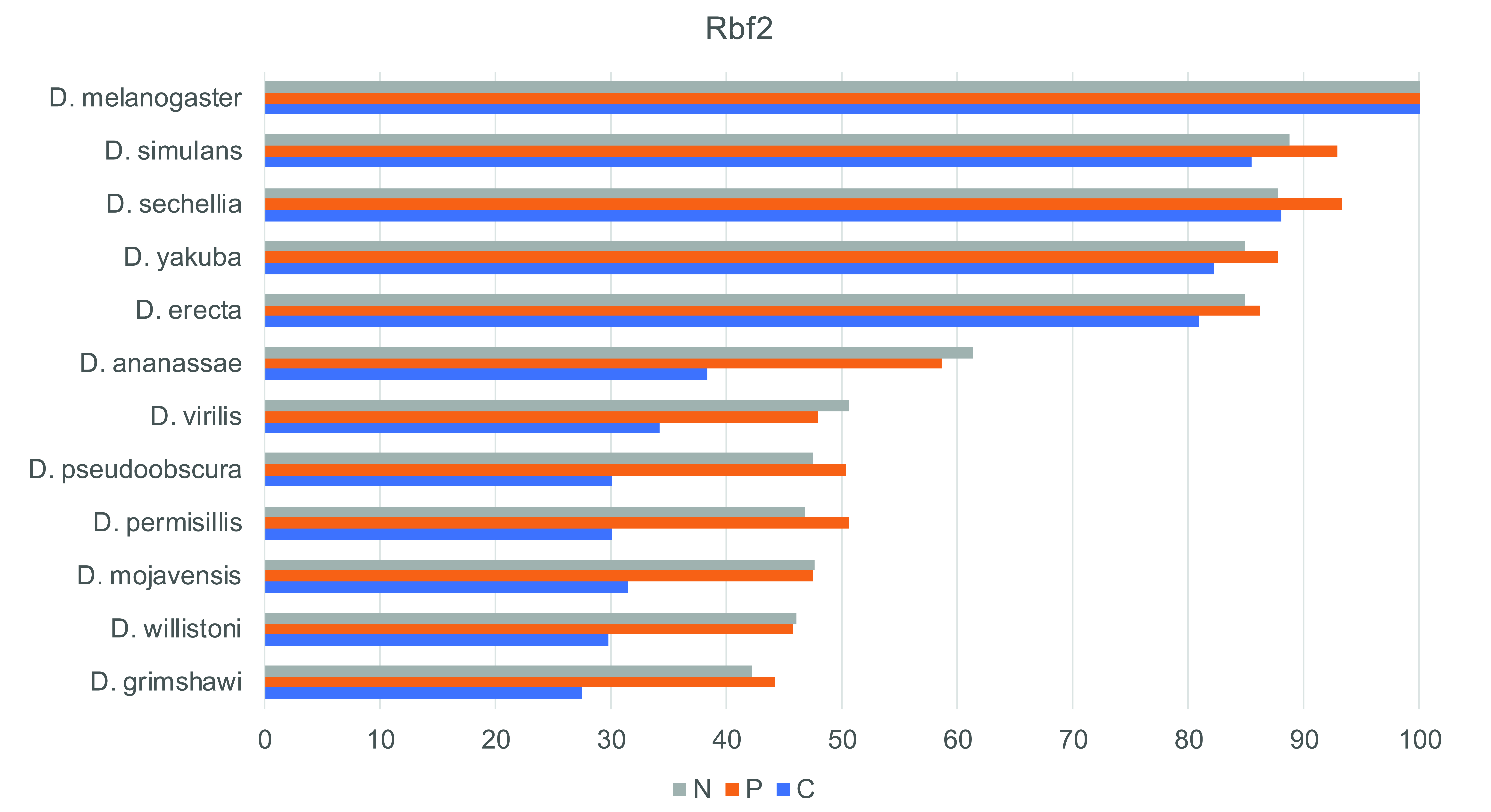

### Supplementary Figure1C

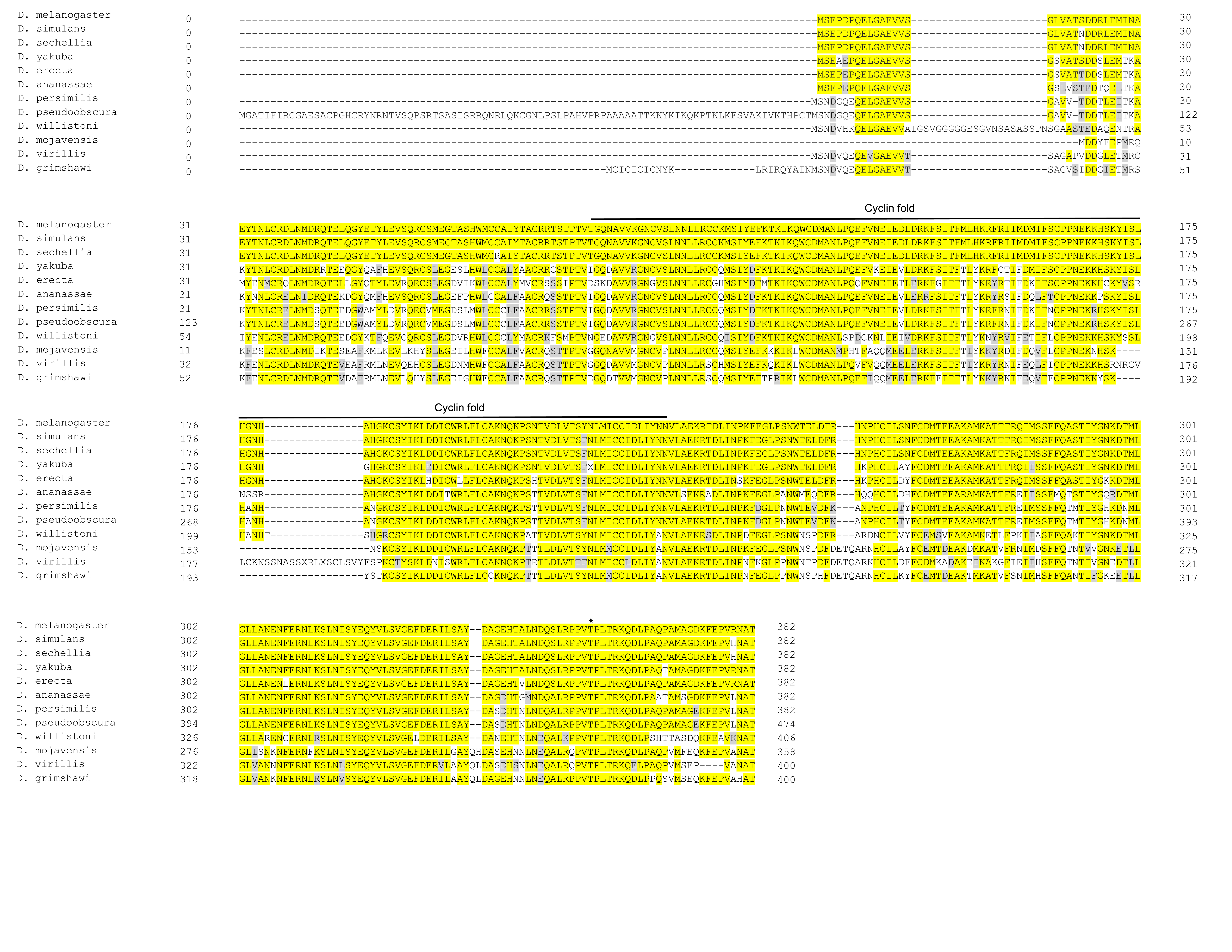

### Supplementary Figure1D

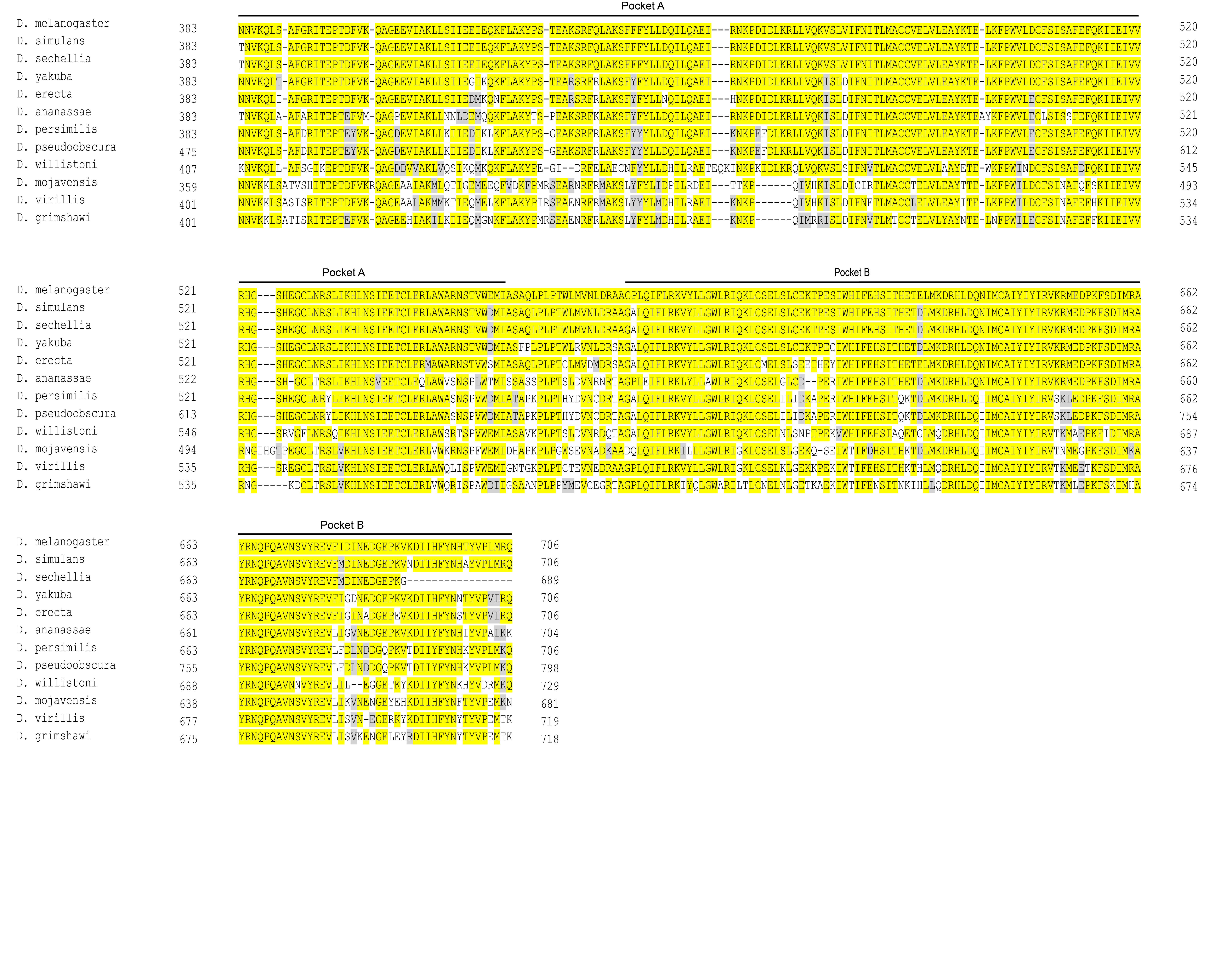

### Supplementary Figure1E

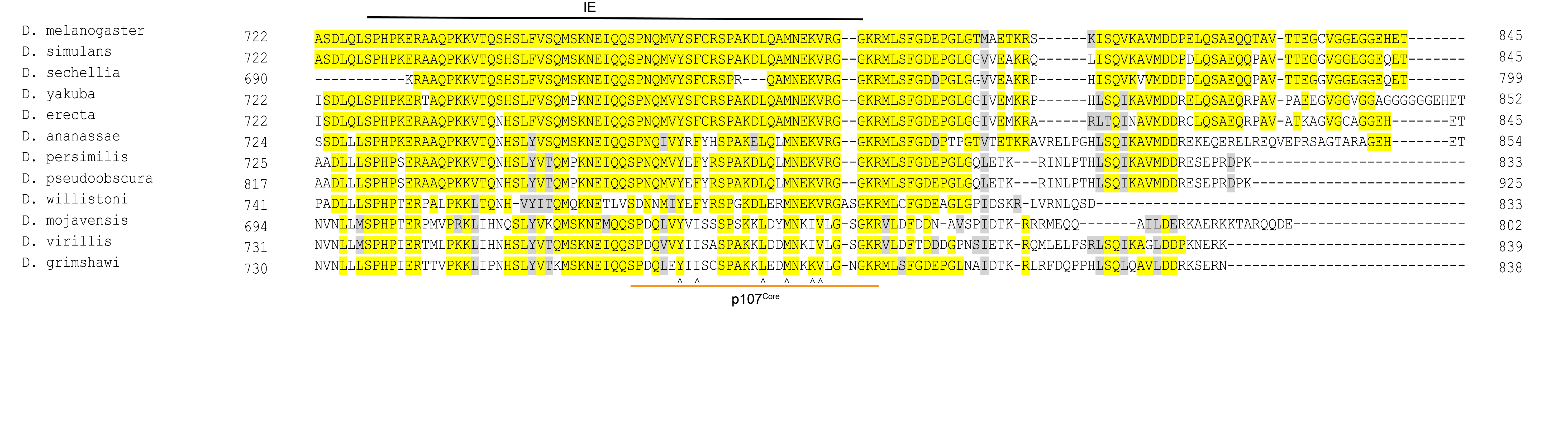

### Supplementary Figure1F

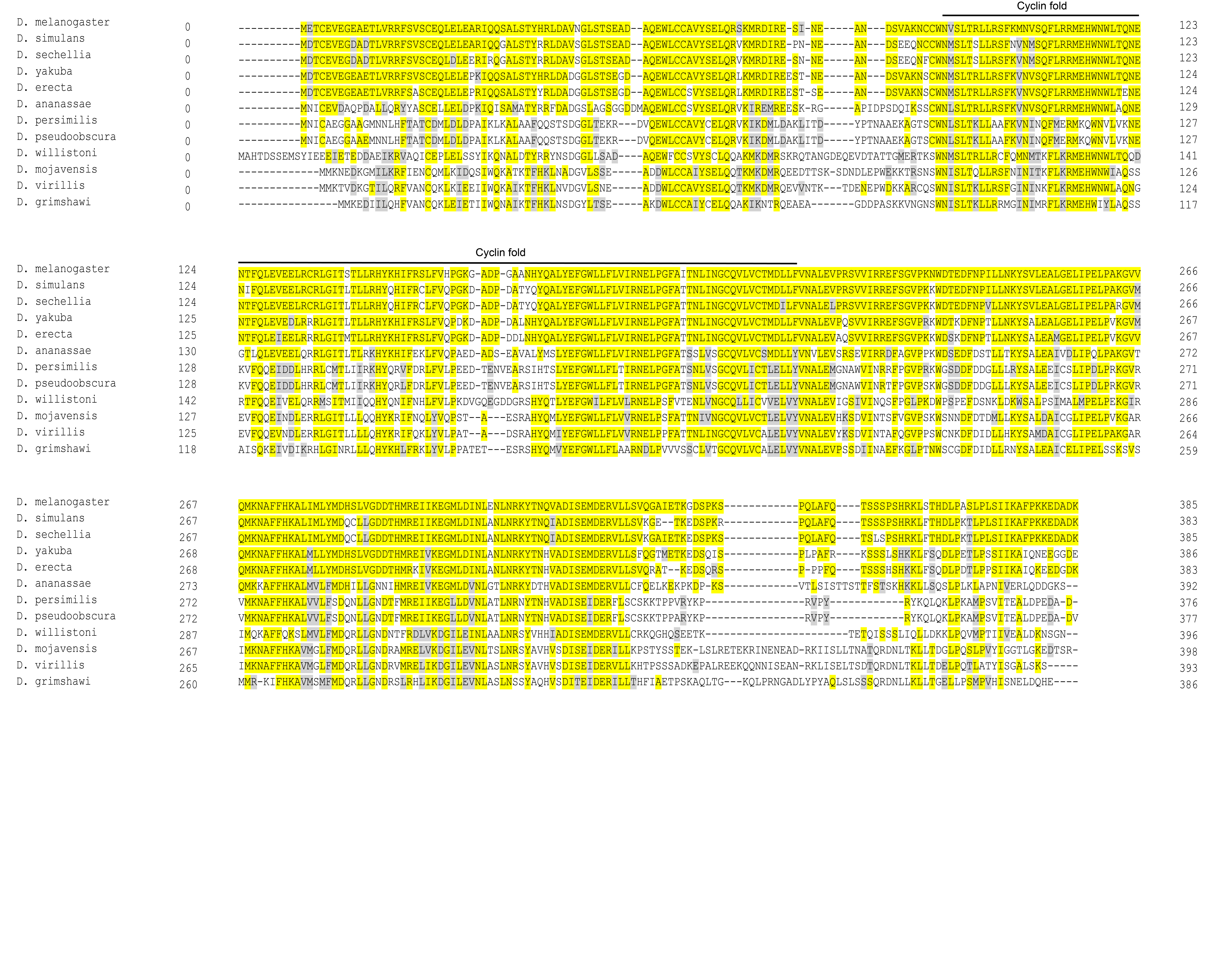

### Supplementary Figure1G

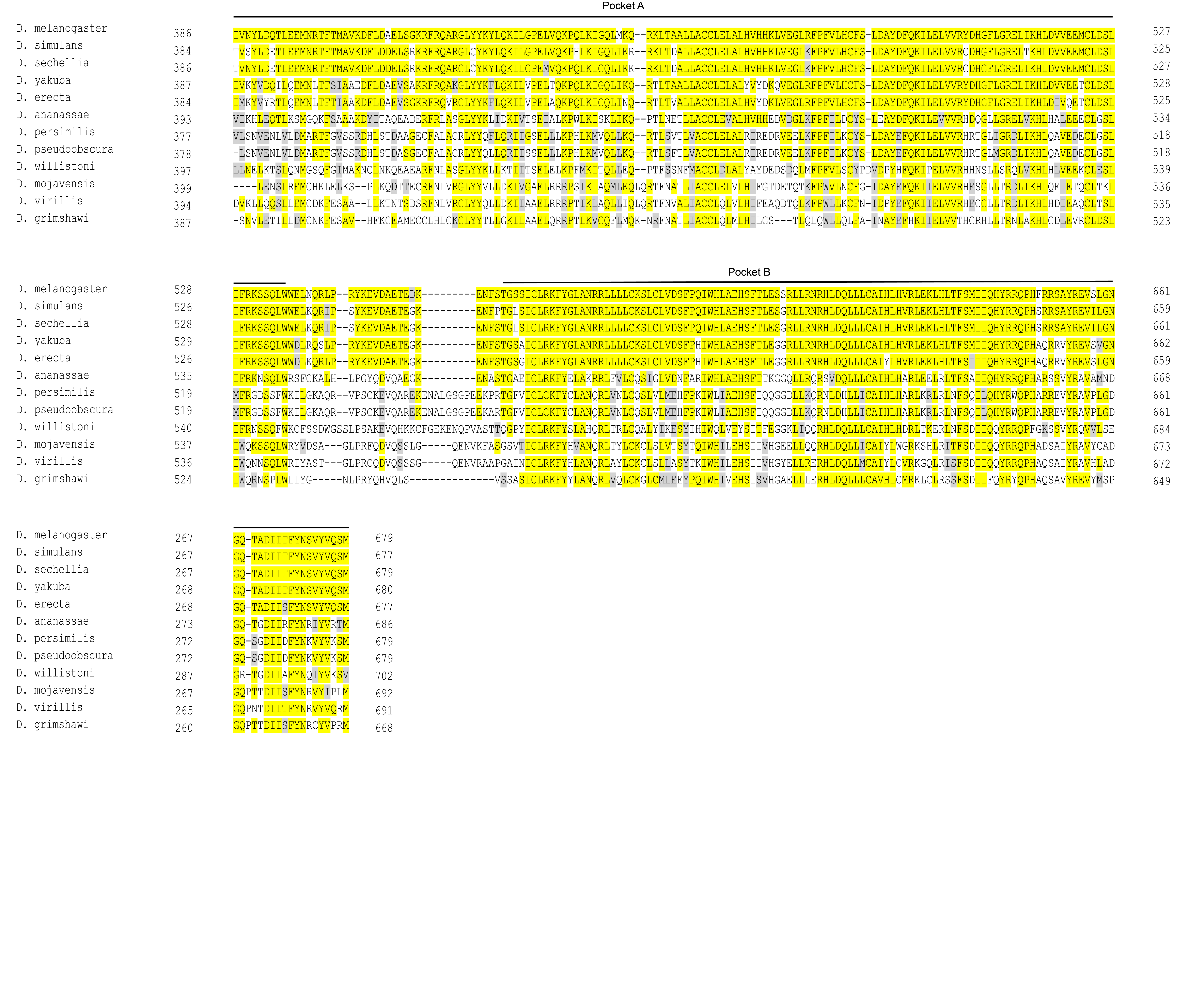

### Supplementary Figure1H

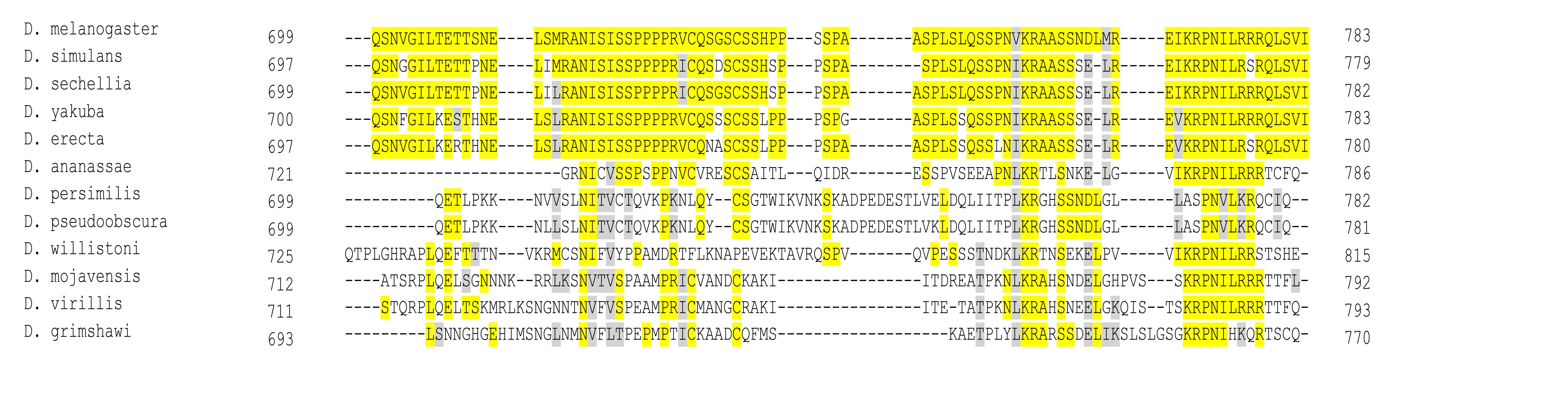

### Supplementary Figure1I

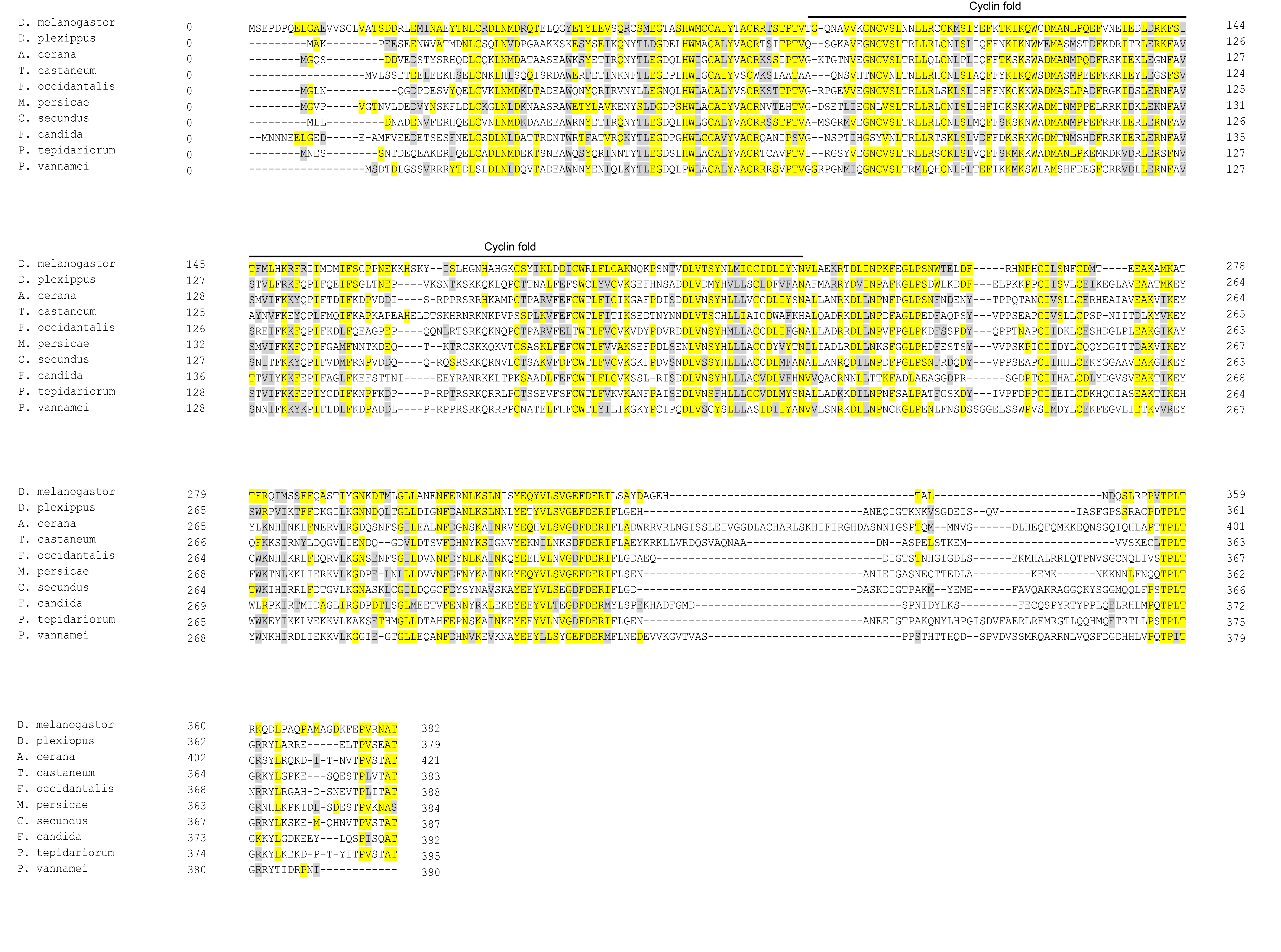

### Supplementary Figure1J

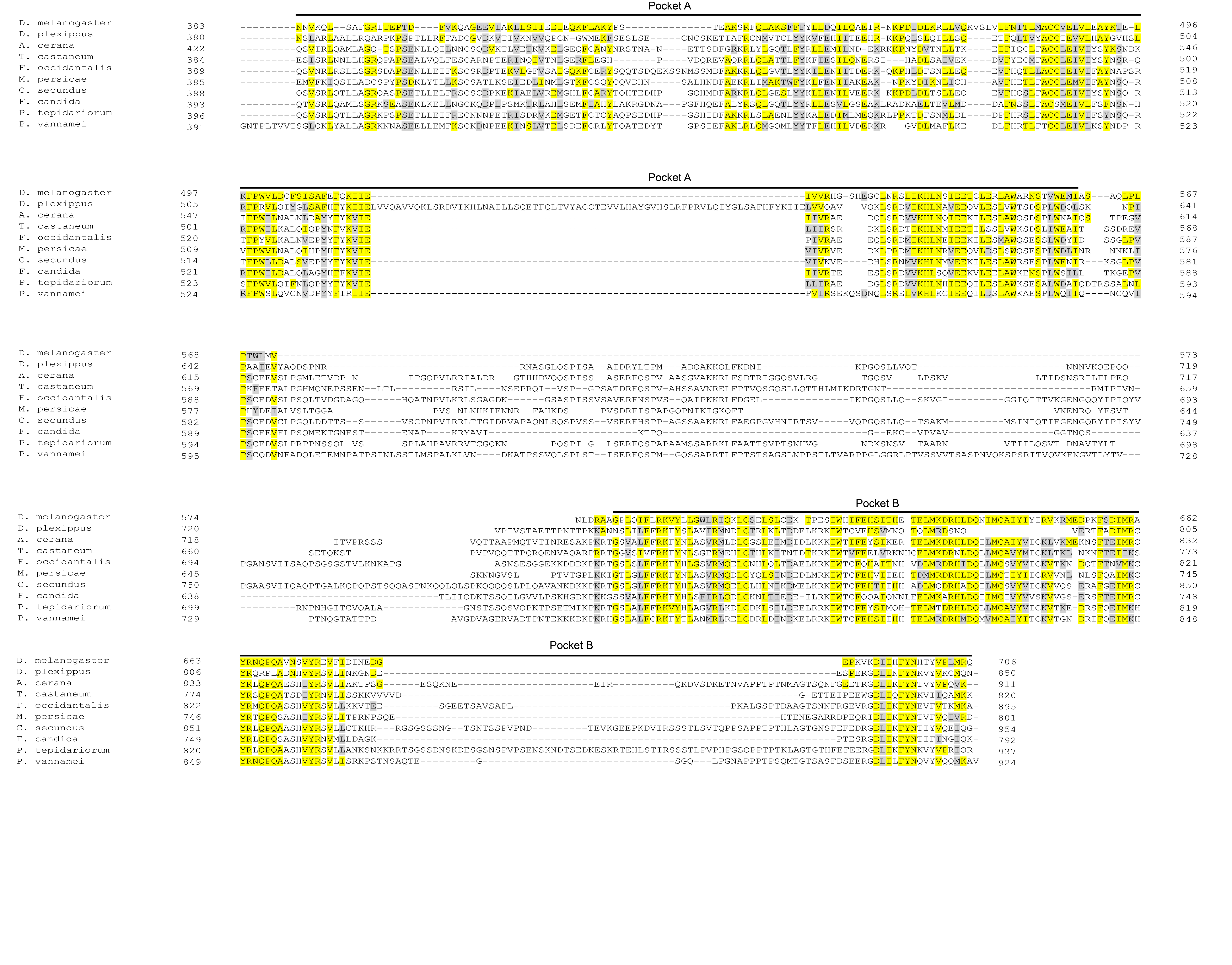

### Supplementary Figure1K

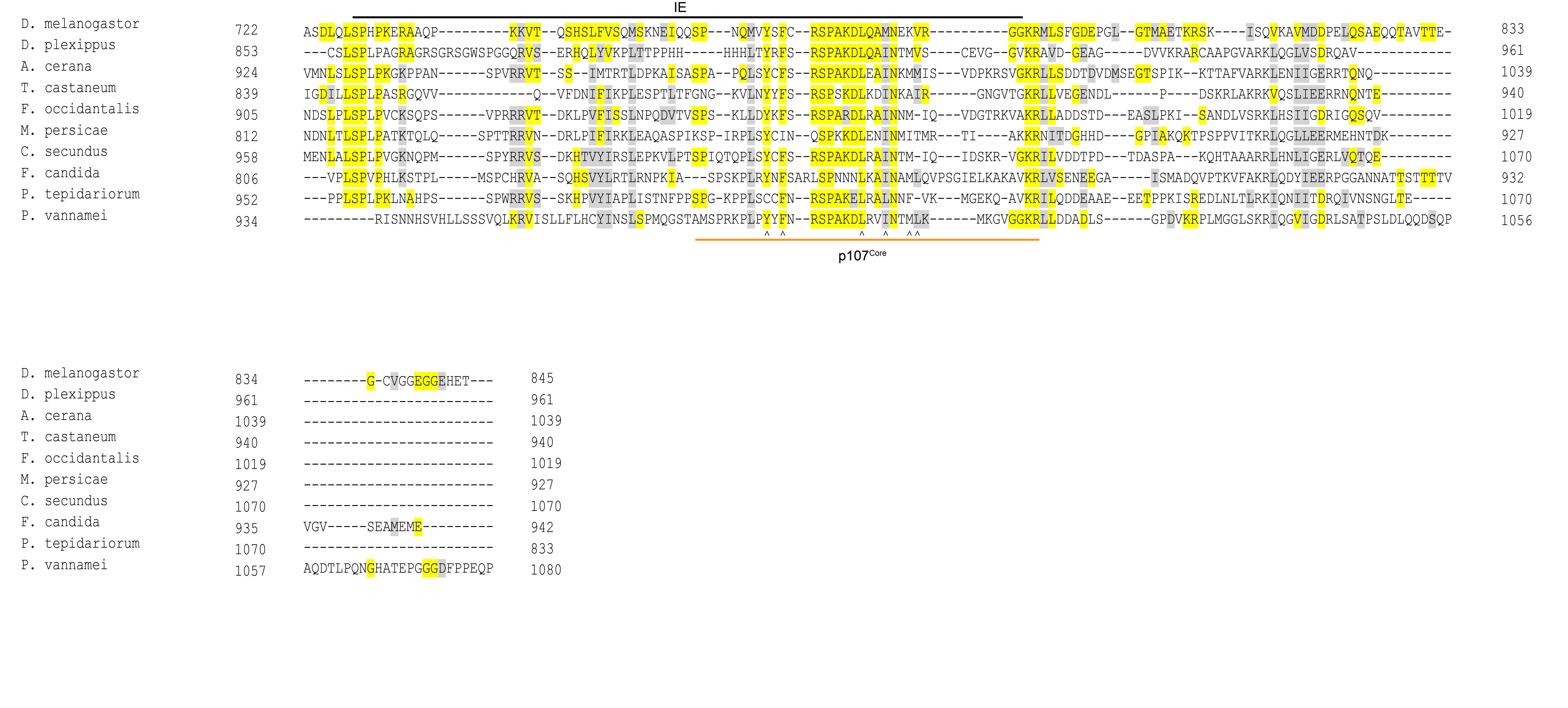

### Supplementary Figure2A

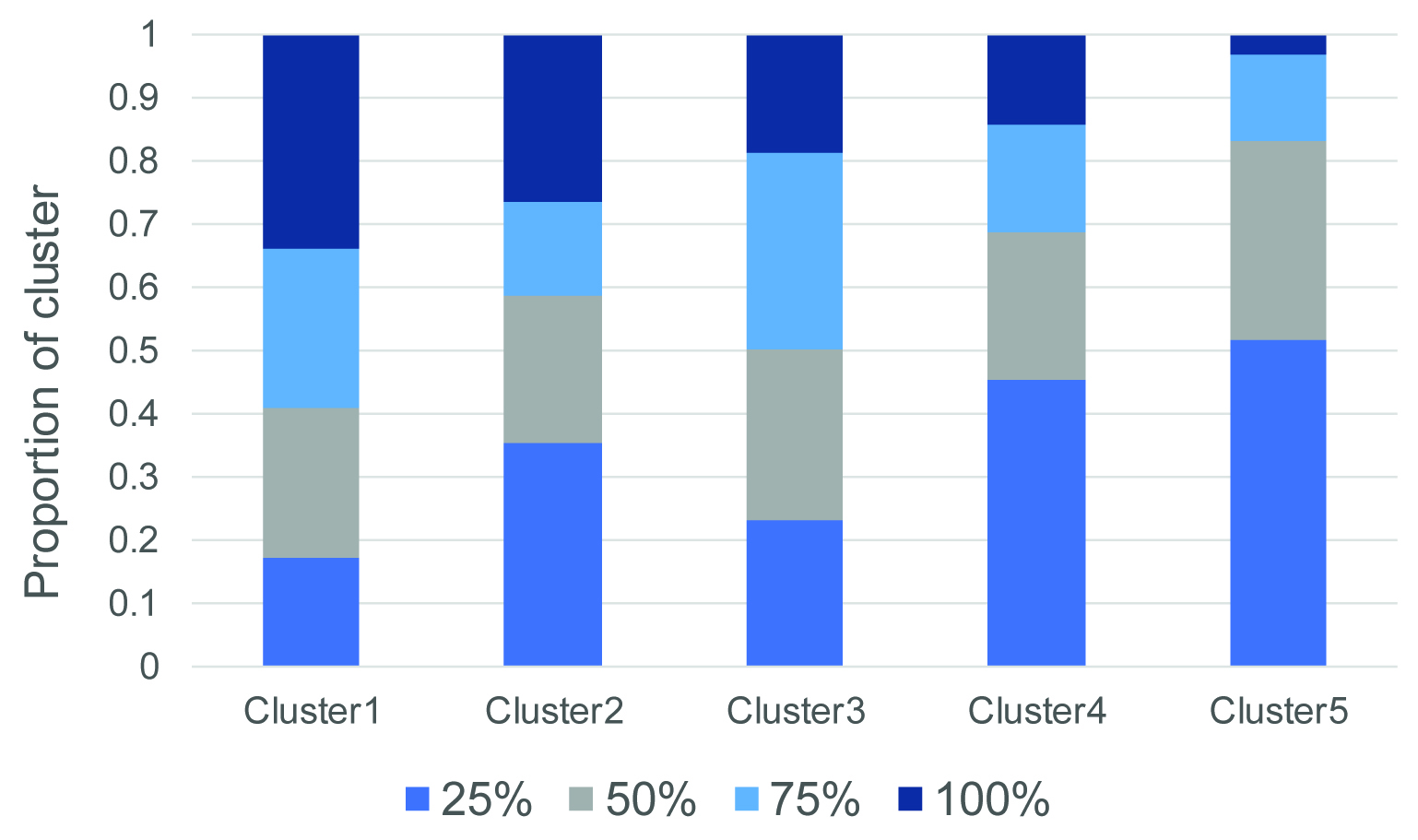

### Supplementary Figure3A

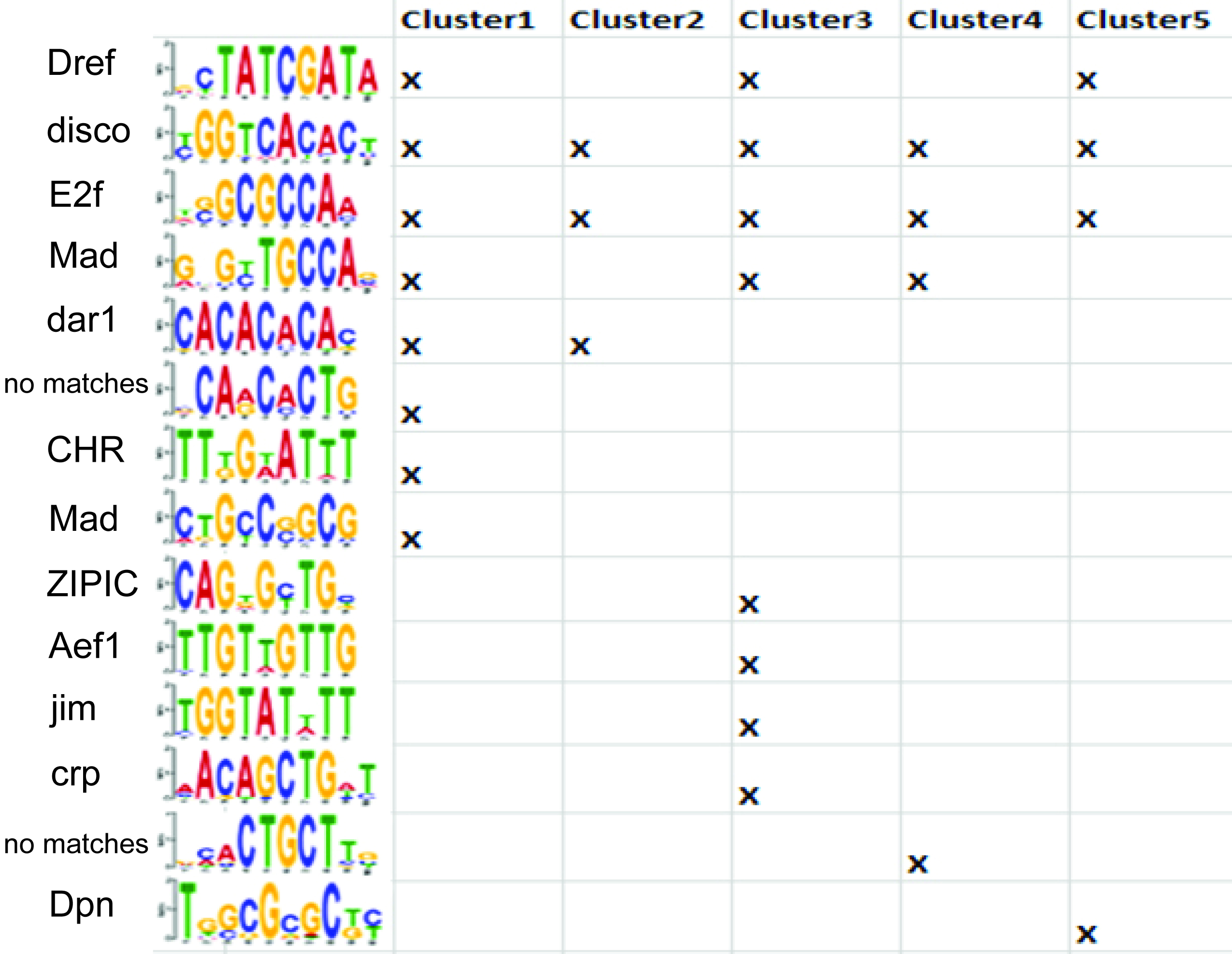

### Supplementary Figure4

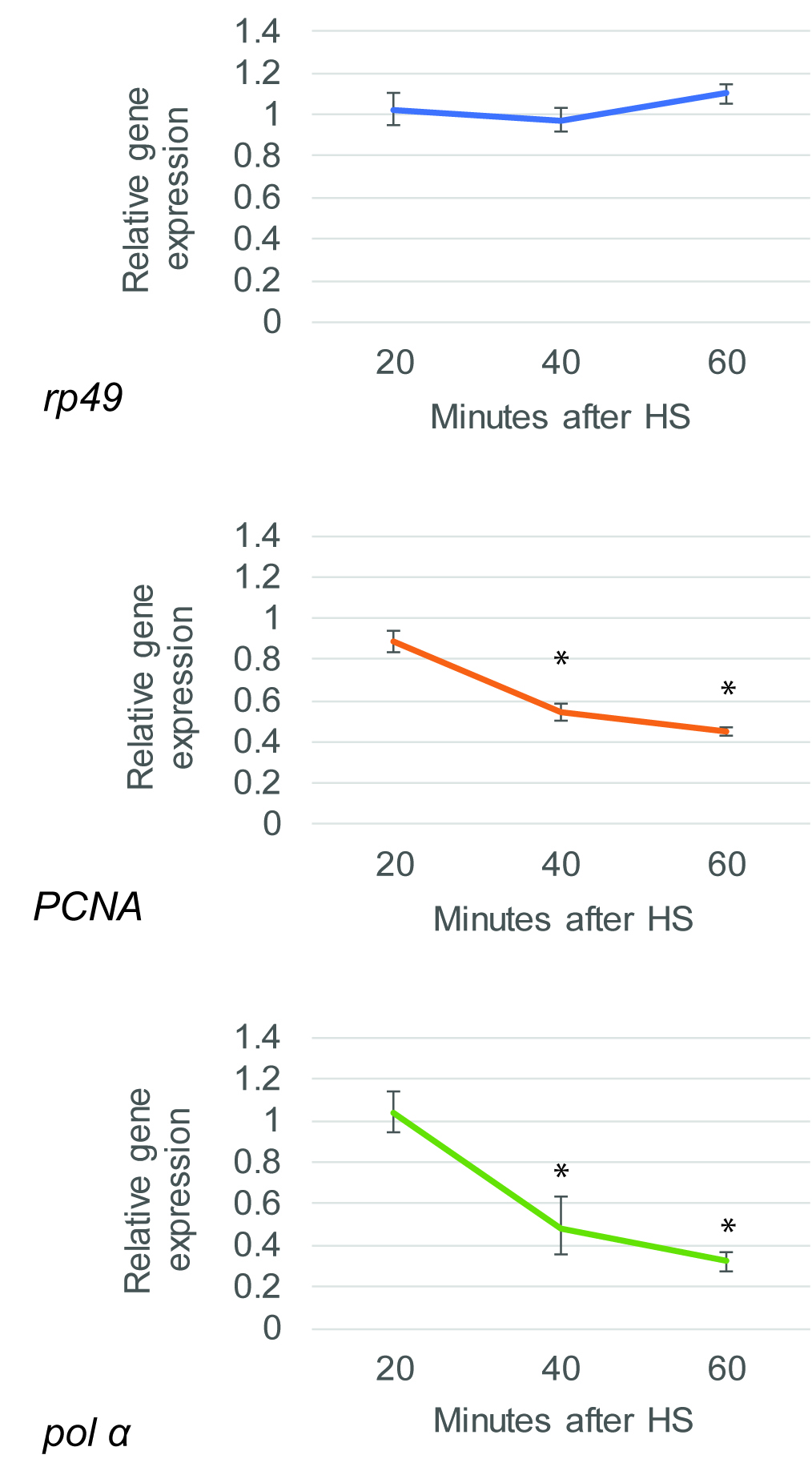

### Supplementary Table2A

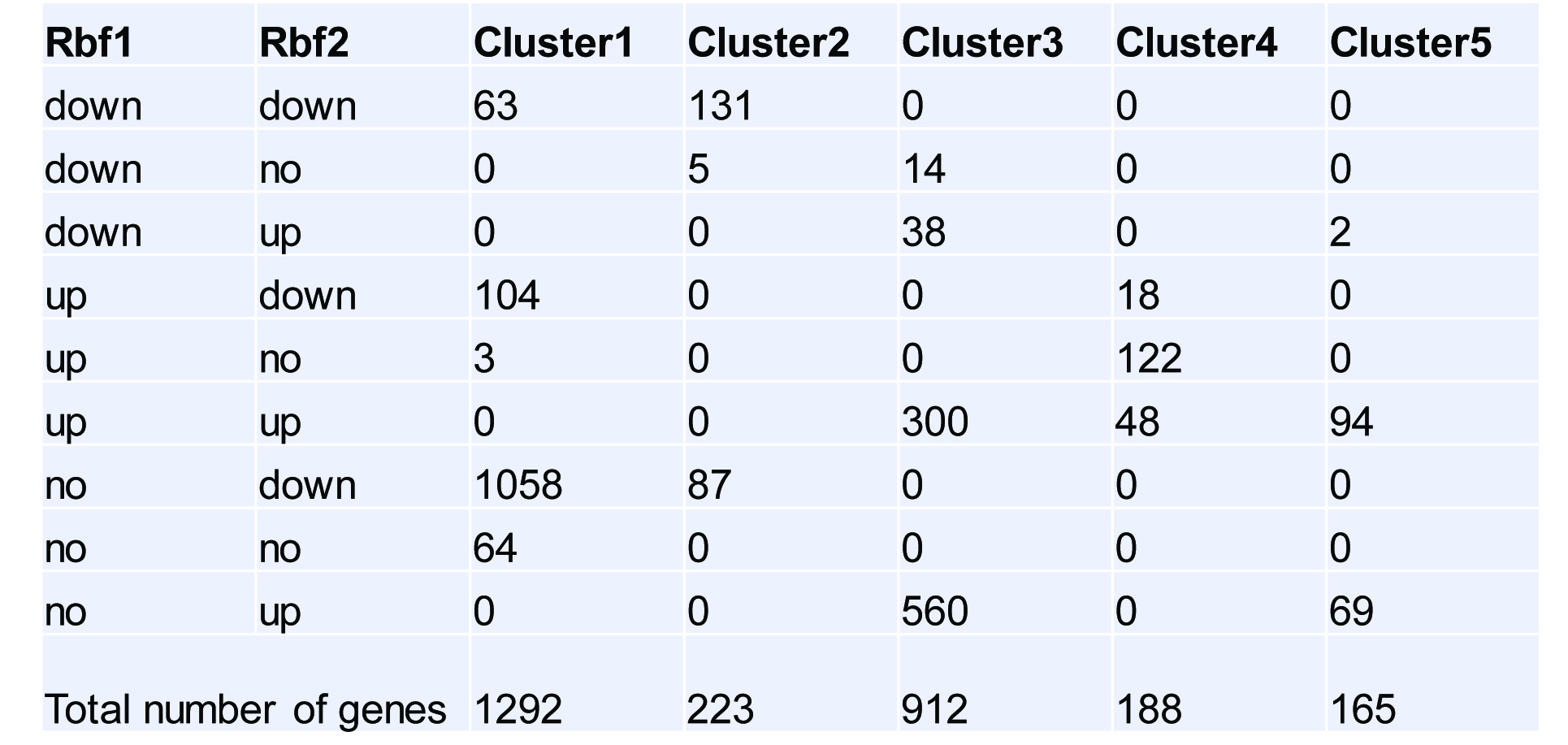

### Supplementary Table2C

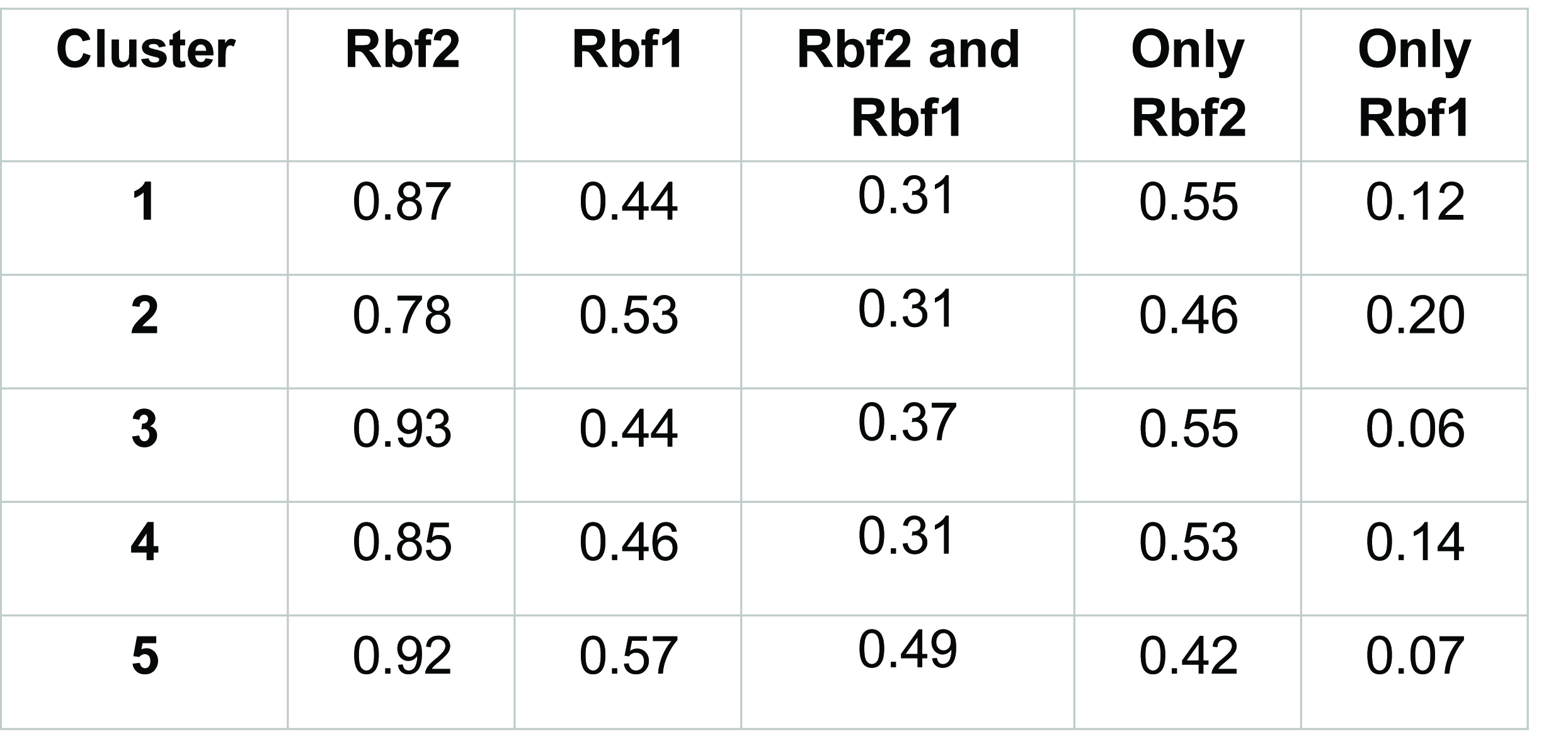

### Supplementary Table3A

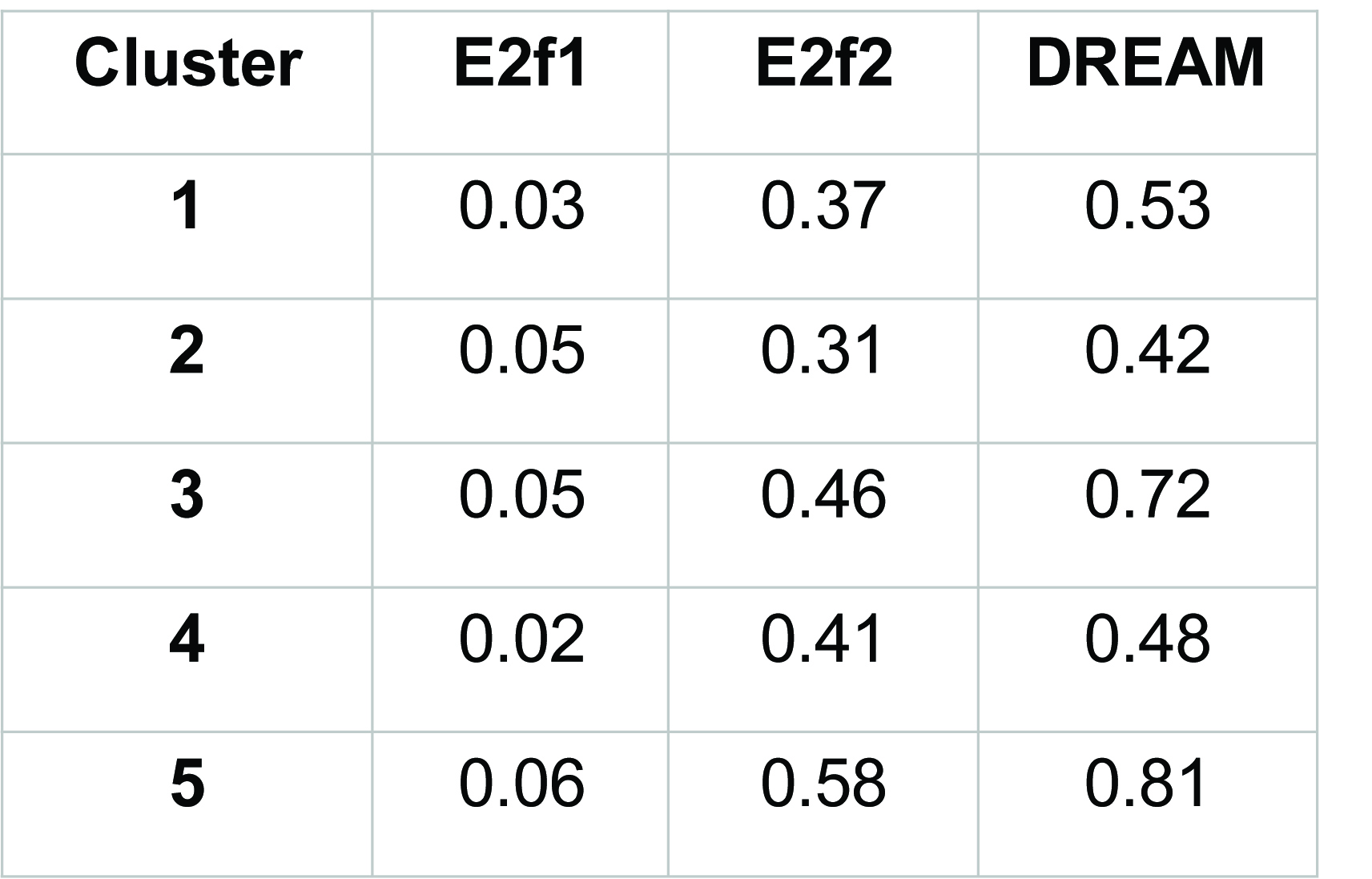
